# Self-delivering CRISPR RNAs for AAV Co-delivery and Genome Editing *in vivo*

**DOI:** 10.1101/2023.03.20.533459

**Authors:** Han Zhang, Karen Kelly, Jonathan Lee, Dimas Echeverria, David Cooper, Rebecca Panwala, Zexiang Chen, Nicholas Gaston, Gregory A. Newby, Jun Xie, David R. Liu, Guangping Gao, Scot A. Wolfe, Anastasia Khvorova, Jonathan K. Watts, Erik J. Sontheimer

## Abstract

Guide RNAs offer programmability for CRISPR-Cas9 genome editing but also add challenges for delivery. Chemical modification, which has been key to the success of oligonucleotide therapeutics, can enhance the stability, distribution, cellular uptake, and safety of nucleic acids. Previously, we engineered heavily and fully modified SpyCas9 crRNA and tracrRNA, which showed enhanced stability and retained activity when delivered to cultured cells in the form of the ribonucleoprotein complex. In this study, we report that a short, fully stabilized oligonucleotide (a “protecting oligo”), which can be displaced by tracrRNA annealing, can significantly enhance the potency and stability of a heavily modified crRNA. Furthermore, protecting oligos allow various bioconjugates to be appended, thereby improving cellular uptake and biodistribution of crRNA *in vivo*. Finally, we achieved *in vivo* genome editing in adult mouse liver and central nervous system via co-delivery of unformulated, chemically modified crRNAs with protecting oligos and AAV vectors that express tracrRNA and either SpyCas9 or a base editor derivative. Our proof-of-concept establishment of AAV/crRNA co-delivery offers a route towards transient editing activity, target multiplexing, guide redosing, and vector inactivation.

## Introduction

Rapidly evolving CRISPR-Cas genome editing tools have revolutionized biomedical research, and promise new routes to therapeutics. Currently, most of the widely applied genome editing tools are based on a Cas9 protein from *Streptococcus pyogenes* (SpyCas9) (1–3). The natural effector complex consists of SpyCas9 bound to a CRISPR RNA (crRNA) and a trans-activating crRNA (tracrRNA) (4–7). The guide sequence within the crRNA can direct the effector complex to a desired genomic locus followed by a specific protospacer adjacent motif (PAM), and the Cas9 nuclease can generate a double-strand DNA (dsDNA) break. When applied to genome editing in eukaryotic cells, non-homologous-end-joining (NHEJ) can induce indels that result in gene disruption, while homology-directed repair (HDR) can generate precise edits (2). HDR-independent precision genome editing tools have also been developed by fusing functional domains to Cas9 nickase (nCas9) mutants. For example, base editors consist of nucleobase deaminase enzymes fused to nCas9 and can generate single-base changes within the targeted genomic locus without requiring dsDNA breaks (8–12). More recently, prime editors were developed by fusing engineered reverse transcriptase (RT) to nCas9, which offers even more versatility and can make virtually any substitution, small insertion, and deletion within the genome (3, 13–15).

The programmability and versatility of CRISPR genome editing tools derive from their RNA-guided nature. However, an RNA guide also adds challenges for *in vivo* delivery. Unmodified single-stranded RNAs (ssRNA) are extremely vulnerable to nuclease degradation and are rapidly degraded in cells and biological fluids such as serum (16, 17). Chemical modification, which has been key to the success of oligonucleotide therapeutics, can enhance the stability, distribution, cellular uptake, and safety of nucleic acids (16–27). Indeed, most of the clinically advanced and FDA-approved antisense oligonucleotides (ASOs) and short interfering RNAs (siRNAs) are fully chemically modified and stabilized for delivery without lipid nanoparticle (LNP) formulation (16, 17, 28–33). Complete chemical modification enables efficient delivery and robust *in vivo* efficacy in multiple tissues including the liver as well as the central nervous system (CNS) (16, 18, 34, 35). Likewise, we reasoned that extensive chemical modification of CRISPR guide RNAs could facilitate delivery *in vivo* and potentially achieve self-delivery without LNPs. For example, self-delivering crRNAs could be administered with an effector (and a tracrRNA, if needed) expressed from a viral vector such as adeno-associated virus (AAV), which is FDA-approved and has been shown to efficiently deliver genome editing agents in a wide range of tissues including liver and CNS (15, 36–39). Separating guide delivery from effector-encoding viral vectors has the potential to limit off-target editing (because the co-delivered guide is only transiently present rather than continuously produced), facilitate multiplexed targeting, increase spatiotemporal control over editing events, enable guide RNA redosing (if needed), and provide a route to inactivate the AAV vector once the desired editing is achieved (25).

Previously, using SpyCas9, we have engineered heavily and fully chemically modified crRNAs and tracrRNAs that exhibit increased stability while retaining editing activity when delivered to cultured cells as a ribonucleoprotein (RNP) complex (23). In this study, we designed fully chemically stabilized protecting oligos (P.O.s) that significantly enhanced the stability and potency of a heavily modified crRNA. Furthermore, when linked with bioconjugates, the P.O.s can boost cellular uptake of the crRNA, acting as a delivery agent without interfering with Cas9 activity. Moreover, we show that the chemically modified crRNAs coupled with P.O.s can generate editing by passive, LNP-independent uptake in cultured cells that stably express effector protein and tracrRNA. Finally, we show that unformulated, chemically modified crRNAs co-delivered with tracrRNA- and effector-expressing AAV vectors support genome editing *in vivo* in adult mice.

## Materials and Methods

### Molecular cloning

The U1a-SpyCas9-miniU6-tracrRNA lentivector used for generating the stable HEK293T-SpyCas9-tracrRNA-TLR-MCV1 reporter cell line shown in **Figure 1a** was cloned by Gibson assembly. The Addgene plasmid #52962 was digested with NotI and BamH1 restriction enzymes, and the lentivector backbone was Gibson-assembled with a miniU6-tracrRNA gene block and U1a-SpyCas9 amplified by PCR from Addgene plasmid #121507. Similarly, the dGFP reporter plasmid was cloned into a lentivector for generating the HEK29T-SpyCas9-ABE-tracrRNA-dGFP reporter cell line shown in **Figure 1a**. First, the Addgene plasmid #52962 was digested with NotI and BamH1 restriction enzymes. Then, the dGFP reporter plasmid was digested with EcoRI and MluI restriction enzymes, followed by Gibson assembly with bridging double-strand DNA (dsDNA) containing homology sequences. A PiggyBac transposon plasmid (Addgene plasmid #159095) was used to integrate the SpyCas9-ABE-tracrRNA expression gene in the HEK293T-SpyCas9-ABE-tracrRNA-dGFP reporter and Hepa1-6-SpyCas9-ABE-tracrRNA stable cell lines shown in **Figure 1a**. Addgene plasmid #159095 was digested with SalI and EcoRV, and the backbone was Gibson-assembled with the CMV enhancer and CbH promoter, SpyCas9-ABE8e, U6-tracrRNA, and hPGK-PuroR. The plasmid for packaging AAV to express tracrRNA (**Figure 4, 5**) was cloned by swapping the sgRNA with tracrRNA in Addgene Plasmid #121508 by overlapping PCR, and removing the EGFP-expression cassette by SpeI and XbaI restriction enzyme digestion, followed by Gibson assembly using a bridging dsDNA. The plasmid for packaging AAV to express SpyCas9 was Addgene Plasmid #121507. Plasmids for packaging AAV to express intein-split SpyCas9-ABE8e and tracrRNA (**Figure 5**) were generated by swapping the sgRNA with tracrRNA sequence using overlapping PCR and restriction cloning. Sequences of original plasmids described in this paper can be found in the **Supplementary note** and will be deposited to Addgene.

**Figure 1.**
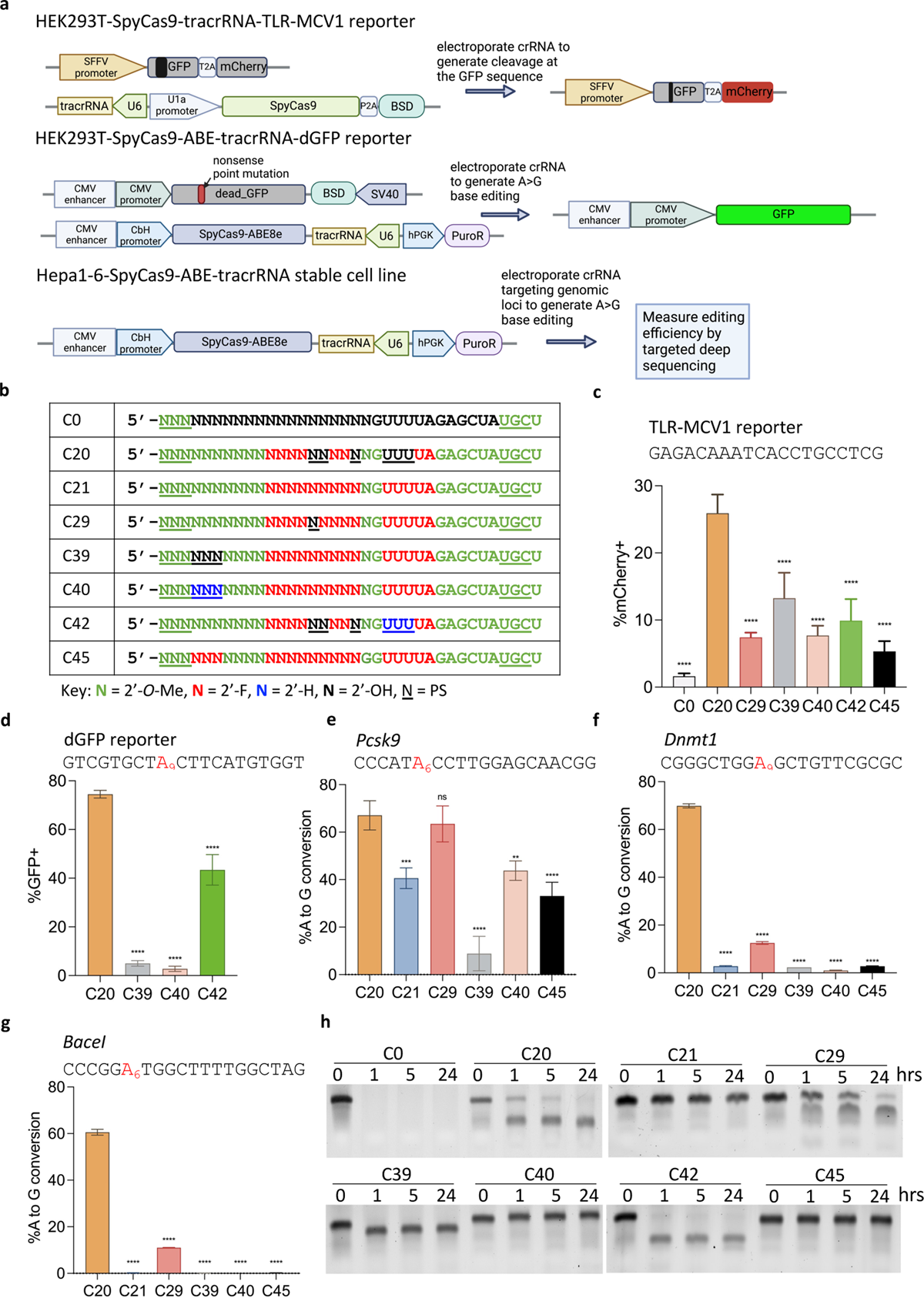
Stable cell lines established and activities of the previously engineered, chemically modified crRNA. a). Schematic representation of stable cell lines for screening the activities of different chemical modifications of crRNA. Top panel: a stable HEK293T cell line that expresses SpyCas9 nuclease, tracrRNA, and the TLR-MCV1 traffic light reporter. Electroporation of crRNAs targeting the broken GFP sequence will generate frameshift mutations that place the mCherry coding sequence in-frame, allowing the editing efficiency to be estimated by mCherry+ quantification via FACS. Middle panel: a stable HEK293T cell line that expresses the adenine base editor SpyCas9-ABE8e, the tracrRNA, and a fluorescent reporter gene that consists of a G-to-A nonsense mutation within the GFP coding sequence [dead_GFP (dGFP)]. Electroporation of crRNA will restore GFP fluorescence by an A-to-G edit that reverses the mutation, and the editing efficiency can then be measured by GFP+ quantification via FACS. Bottom panel: a stable mouse Hepa1-6 cell line that expresses SpyCas9-ABE8e and tracrRNA. Electroporation of crRNA targeting endogenous genomic loci will generate A-to-G edits that can then be measured by targeted amplicon deep sequencing. b). The modification patterns of the previously engineered, heavily, or fully modified crRNAs used in panels c-g. Modified crRNAs targeting different sequences were delivered by electroporating 100 pmol crRNA into 5 x 10^4^ cells in different stable cell lines: c). HEK293T-SpyCas9-tracrRNA-TLR-MCV1 reporter; d). HEK293T-SpyCas9-ABE-tracrRNA-dGFP reporter; e-g), Hepa1-6-SpyCas9-ABE-tracrRNA stable cell line for three endogenous loci, as indicated. The editing efficiencies were quantified by either FACS analysis (c-d) or targeted amplicon deep sequencing (e-g) (n = 3 biological replicates). Data represent mean ± SD; **, P < 0.01; ***, P < 0.001; ****, P < 0.0001 (one-way ANOVA; C20 was used as control in multiple comparisons). h). Stability of partially and fully-modified crRNAs shown in 1b in 10% non-heat-inactivated FBS at 37°C. Digested fragments were resolved by 10% urea-denaturing polyacrylamide gel electrophoresis (PAGE) followed by SYBR Gold staining.

### Stable cell lines

The SpyCas9-tracrRNA transgene in the HEK293T-SpyCas9-tracrRNA-TLR-MCV1 reporter cell line, and the dGFP reporter transgene in the HEK293T-SpyCas9-ABE-tracrRNA-dGFP reporter cell line, were integrated into the genome of HEK293T-TLR-MCV1 reporter cells(40) and HEK293T cells, respectively, by lentiviral transduction. HEK293-F cells (ThermoFisher R79007) were transfected with the transfer plasmid and the packaging plasmids (Addgene plasmids #12263 and #8454) using TransIT-LT1 transfection reagent (Mirus MIR 2304). Two days later, the media containing lentivirus was collected and filtered through a 0.45 µm filter (Cytiva 6780-25040) to remove cell debris. HEK293T-TLR-MCV1 cells were transduced with lentivirus encoding SpyCas9-tracrRNA, and the HEK293T cells were transduced with lentivirus encoding the dGFP reporter in the presence of 8 µg/ml polybrene (Millipore Sigma TR-1003-G). Three days after transduction, the media was removed, and fresh media containing 5 µg/ml Blasticidin (Gibco R21001) was added to both cell lines. Ten days post-selection, cells were collected, and single-cell clones were established by serial dilution in 96-well plates. The SpyCas9-ABE-tracRNA transgene was integrated by PiggyBac transposon into the HEK293T-dGFP reporter cell line to generate HEK293T-SpyCas9-tracrRNA-dGFP reporter cell line, or into Hepa1-6 cell line to generate Hepa1-6-SpyCas9-ABE-tracrRNA stable cell line. Briefly, the SpyCas9-ABE-tracrRNA PiggyBac transposon plasmid and PiggyBac transposase expression plasmid (System Biosciences PB210PA-1) were mixed in a 2:1 molar ratio and transfected into the HEK293T-dGFP reporter cell line using Lipofectamine 3000 (ThermoFisher L3000015), or electroporated into Hepa1-6 cells using a Neon Transfection System 10 µl kit (ThermoFisher MPK 1096) with the following electroporation parameters: pulse voltage (1230 v), pulse width (20 ms), pulse number (3). Four days after transfection or electroporation, fresh media containing 2.5 µg/ml Puromycin (ThermoFisher A1113802) was added to both cell lines. Seven days post-selection, cells were collected, and single-cell clones were established by serial dilution in 96-well plates.

### Cell culture

HEK293T reporter cell lines and the Hepa1-6 stable cell line were cultured in Dulbecco’s Modified Eagle Media (DMEM, Genesee Scientific 25-500) supplemented with 10% Fetal Bovine Serum (FBS, Gibco 26140079). All cells were incubated in a 37 °C incubator with 5% CO2.

### Oligonucleotide Synthesis

Oligonucleotides were synthesized by phosphoramidite solid-phase synthesis on a Dr. Oligo 48 (Biolytic, Fremont, CA). All phosphoramidites with standard protecting groups were purchased from ChemGenes (Wilmington, MA). Phosphoramidites were prepared at 0.1 M in anhydrous acetonitrile (ACN), except for 2’-*O*-methyl-uridine dissolved in anhydrous ACN containing 15% dimethylformamide. 5-(Benzylthio)-1H-tetrazole (BTT) was used as the activator at 0.25 M, and the coupling time for all phosphoramidites was 4 minutes. Detritylations were performed using 3% trichloroacetic acid in dichloromethane. Capping reagents used were CAP A (20% n-methylimidazole in ACN) and CAP B (20% acetic anhydride and 30% 2, 6-lutidine in ACN). Reagents for capping and detritylation were purchased from American International Chemical LLC (AIC), Westborough, MA. Phosphite oxidation to convert to phosphate was performed with 0.05 M iodine in pyridine-H2O (9:1, v/v) (AIC) or phosphorothioate with a solution of 0.1 M of 3 [(dimethylaminomethylene)amino]-3H-1, 2, 4-dithiazole-5-thione (DDTT) in pyridine (ChemGenes) for 4 minutes. Unconjugated oligonucleotides were synthesized on 500 Å long-chain alkyl amine (LCAA) controlled pore glass (CPG) functionalized with Unylinker terminus (ChemGenes). Cholesterol-conjugated oligonucleotides were synthesized on a 500Å LCAA-CPG support, where the cholesterol moiety is bound through a tetra-ethyleneglycol (TEG) linker (Chemgenes, Wilmington, MA). GalNAc conjugated oligonucleotides were grown on a 500Å LCAA-CPG where the trivalent GalNAc cluster was linked to the oligonucleotide through an aminopropanediol linker. TEG was used as the proximal spacers for each of the GalNac units. DHA-conjugated oligonucleotides were synthesized on a 500Å LCAA-CPG where the DHA is bound to the oligonucleotide via a C7-linker. Both GalNAc and DHA CPGs were purchased from Hongene Biotech, Union City, CA.

### Deprotection and Purification of Oligonucleotides

Synthesis columns containing oligonucleotides were treated previously with 10% diethylamine (Fisher) in ACN on the synthesizer. Oligonucleotides were cleaved and base-protecting groups were removed with a solution of 1:1 40% methylamine in water/30% ammonium hydroxide for 2 hours at room temperature. Cleaved and deprotected oligos were fully dried under a vacuum. Oligonucleotides containing 2’TBDMS protecting groups were dissolved in 115 µL DMSO (Sigma-Aldrich) at 65 °C. Triethylamine 60 µL (Sigma-Aldrich) followed by triethylamine-trihydrofluoride 75 µL (Sigma-Aldrich) was added, and the whole solution was incubated for 2.5 hours at 65°C. Deprotected oligonucleotides were cooled and precipitated in a solution of 0.1 M sodium acetate in isopropanol (Sigma Aldrich). After centrifugation, the supernatants were discarded, and the pellets were dried under a vacuum. Dried oligonucleotides were dissolved in 400 µL RNase-free water and desalted using Amicon Ultra 0.5 mL 3K filter tubes (Millipore, Billerica, MA USA), followed by three RNase-free water wash rounds for 15 min at 14000 x g. Finally, desalted oligonucleotides were extracted and diluted in RNase-free water.

### Quantification and LC-MS Analysis of Oligonucleotides

Absorbances at 260 nm were obtained using a Nanodrop 1000 series. Extinction coefficients were calculated for each sequence, and concentrations were determined following the Beer-Lambert equation. The identities of oligonucleotides were verified by LC-MS analysis on an Agilent 6530 accurate mass Q-TOF using the following conditions: buffer A: 100 mM 1, 1, 1, 3, 3, 3-hexafluoroisopropanol (HFIP) and 9 mM triethylamine (TEA) in LC-MS grade water; buffer B:100 mM HFIP and 9 mM TEA in LC-MS grade methanol; column, Agilent AdvanceBio oligonucleotides C18; linear gradient 5-100% B in 5 minutes was used for all oligonucleotides; temperature, 60 °C; flow rate, 0.85 ml/min. LC peaks were monitored at 260 nm. MS parameters: Source, electrospray ionization; ion polarity, negative mode; range, 100–3, 200 m/z; scan rate, 2 spectra/s; capillary voltage, 4, 000; fragmentor, 200 V; gas temp, 325 °C.

### Buffer exchange of oligonucleotide for *in cellulo* and *in vivo* study

Oligonucleotides were first concentrated using Ultra Centrifugal 3K Filter Unit (Amicon UFC500396), then washed three times with 1M Sodium Acetate solution (Sigma 711096) and three times with phosphate buffered saline (PBS), pH 7.4 (Gibco 10010-023). Finally, the oligonucleotides were eluted in PBS, pH 7.4 (Gibco 10010-023), and absorbances at 260 nm were obtained using Nanodrop 1000 to determine the concentrations for *in cellulo* and *in vivo* study.

### CrRNA/protecting oligo (P.O.) annealing and electroporation

Each crRNA was mixed with P.O. at a 1:1 ratio, heated to 95 °C for 5 minutes, then gradually cooled to 15 °C (−0.1 °C per second, with the temperature held for 30 seconds every 10 °C). Electroporation of crRNA was performed using a Neon Transfection System 10 µl kit (ThermoFisher MPK 1096), with the following parameters: HEK293T cells, pulse voltage (1150 v), pulse width (20 ms), pulse number (2); Hepa1-6 cells, pulse voltage (1230 v), pulse width (20 ms), pulse number (3). Cells were cultured for 48 hours before being quantified by flow cytometry.

### Free uptake of crRNA

HEK293T-SpyCas9-ABE-tracrRNA-dGFP reporter cells were seeded at 25% confluency in a 48-well cell culture plate (ThermoFisher 150687) and cultured in DMEM + 10% FBS for 24 hours. Fresh culture media containing DMEM + 3% FBS and various concentrations of crRNA were mixed with the cells by gently pipetting up and down and incubated for an additional 72 hours before being quantified by flow cytometry.

### Fluorescent reporter assay

Cells were trypsinized, harvested into microcentrifuge tubes and centrifuged at 300 x g for 3 mins. Supernatant was removed and the cell pellet was resuspended into 1 x PBS and transferred into flow cytometry tubes. MACSQuant VYB was used for quantification of the percentage of fluorescent-positive cells, and 10, 000 events were counted for FACS analysis. Data was analyzed using Flowjo v10. A representative gating strategy can be found in **Supplementary Figure 1**. **crRNA stability assay.** The crRNAs with or without P.O. were diluted in PBS, pH 7.4 and 10% FBS (ThermoFisher A3160501) and incubated at 37 °C for 0, 1, 4, 24 hours in a thermocycler. At each time point, samples were taken and mixed with an equal volume of 2x RNA gel loading dye (ThermoFisher R0641) and heated to 75 °C for 10 minutes to denature. The digested RNA was further resolved by 10% urea polyacrylamide gel electrophoresis (PAGE) followed by SYBR Gold (Invitrogen S11494) staining.

### AAV production

AAV vector packaging was done at the Viral Vector Core of the Horae Gene Therapy Center at the UMass Chan Medical School as previously described (41). Constructs were packaged in AAV9 capsids, and viral titers were determined by digital droplet PCR and gel electrophoresis, followed by silver staining.

### Stereotactic intrastriatal (IS) injection

All animal study protocols were approved by the Institutional Animal Care and Use Committee (IACUC) at UMass Chan Medical School. The Cas9^+/+^ mice that constitutively express Cas9 was generated by crossing a male mouse of Jackson Lab #027632 with a Cre-expressing female mouse (JAX #008454). The Cas9/mTmG mice were generated by crossing the Cas9^+/+^ mice and the mTmG^+/+^ mice (JAX #007676). For IS injection, 10 - 13 week-old mice were weighed and anesthetized by intraperitoneal injection of a 0.1 mg/kg Fentanyl, 5 mg/kg Midazolam, and 0.25 mg/kg Dexmedetomidine mixture. A total dose of 5 x 10^10^ vg of AAV in 3 µl was administered via unilateral intrastriatal injection (3 µl on the right side) performed as previously described at the following coordinates from bregma: +1.0 mm anterior-posterior (AP), ±2.0 mm mediolateral, and −3.0 mm dorsoventral (35, 42). Once the injection was completed, mice were intraperitoneally injected with 0.5 mg/kg Flumazenil and 5.0 mg/kg Atipamezole, and subcutaneously injected with 0.3 mg/kg Buprenorphine. Two weeks later, the same IS injection was performed to deliver 3 nmol crRNA in 3 µl volume into the AAV-injected site. Mice were euthanized 2 weeks after the second injection.

### Retro-orbital (RO) injection

The *Fah*^PM/PM^ mice were kept on water supplemented with 10 mg/L 2-(2-nitro-4-trifluoromethylbenzoyl)-1, 3-cyclohexanedione (NTBC; Sigma, PHR1731-1G). Mice with more than 20% weight loss were humanely euthanized according to IACUC guidelines. For RO injection, six-week-old female mice were anesthetized by placement inside an approved induction chamber and introduction of Isoflurane at a flow rate of 2-3% and oxygen at 1.5 liters per minute. A total dose of 2 x 10^12^ vg of AAV (1 x 10^12^ vg of each AAV) in 200 µl was injected into the retro-orbital sinus of the mice. Animals were observed for signs of pain or distress and returned to a clean cage when able to remain sternal. Five weeks later, the same RO injections were performed to deliver GalNac-crRNA. When multiple injections of retro-orbital injection were performed, the injection site was alternated between the two eyes.

### Genomic DNA extraction from cultured cells and mouse tissues

For cultured cells, genomic DNAs were extracted 48 hours after electroporation. Briefly, cell culture media were aspirated, and cells were lysed using QuickExtract DNA Extraction Solution (Lucigen) following the manufacturer’s protocols. For mouse tissues, biopsies from each of the five liver lobes were collected into individual tubes. Tissues were homogenized, and genomic DNA was extracted using GenElute Mammalian Genomic DNA Miniprep Kit (Millipore Sigma G1N350).

### Immunohistochemistry (IHC)

A floating section IHC staining protocol (43) was used in **Figure 4c**. Briefly, mice were perfused with PBS and brains were harvested and fixed with 10% neutral buffered formalin (ThermoFisher 5735) for 20 hours. Coronal brain sections (40 µm) were obtained using a vibratome (SCHOTT KL 1500). Brain sections were permeabilized by incubating in PBS, pH 7.4 with 0.3 % Triton X-100 (Sigma T8787), and blocked with 3% hydrogen peroxide (Sigma 216763). The sections were then incubated with blocking solution [1.5% normal goat serum (Vector Laboratories S-1000) + 0.3 % Triton X-100 in PBS, pH 7.4] for 3 hours at room temperature (RT) followed by incubating with a rabbit anti-EGFP antibody (ThermoFisher G10362, 1:1000 dilution) at 4 °C overnight. After washing with PBS, the sections were incubated with a secondary antibody (Vector Laboratories BA-1000) for 10 minutes at RT, then with ABC reagent (Vector Laboratories PK-6100) for 30 minutes at RT. To detect EGFP-positive signals, the sections were stained with 1x DAB (ThermoFisher 34065) for 2 minutes at RT. Finally, the sections were mounted on slides with PBS and 0.5% gelatin (Sigma G1890) in 30% ethanol and dried overnight before imaging.

IHC staining shown in **Figure 5h** used formalin-fixed, paraffin-embedded sections. Briefly, mice were euthanized by CO2 asphyxiation, and livers were fixed with 10% neutral buffered formalin (Epredia 5735), embedded with paraffin, and sectioned at 10 µm. For IHC, liver sections were dewaxed, rehydrated, and stained using an anti-FAH antibody (Abcam ab83770) at 1:400 dilution as described previously (42, 44).

### ImageJ analysis on IHC images

The percentage of EGFP-positive cells was quantified using the ImageJ color deconvolution 2 plugin. A 1 mm^2^ region around the injection site was selected for quantification. Colors were deconvoluted using “H DAB”, and the number of nuclei was counted by inverting the nuclei image color, applying median blur (3), and find maxima (30). The number of EGFP-positive cells was counted by inverting the DAB image color, applying Gaussian blur (2), setting the threshold to remove the background counting, and applying watershed, followed by analyzing particles (200 - infinity). The percentage of EGFP-positive cells = (number of DAB-positive cells/number of nuclei) x 100%.

### Fluorescent imaging

Mouse brains IS-injected with Cy3-labeled crRNAs were perfused with PBS, fixed with 10% neutral buffered formalin for 20 hours, paraffin-embedded, and sectioned to 10 µm. Before imaging, the brain sections were deparaffinized by incubating in xylene for 10 minutes, and mounted with DAPI mounting media (Invitrogen P36962). Images were taken under a Leica fluorescent microscope. The same laser intensity and exposure settings were applied to all images for comparisons between groups.

### Targeted amplicon deep sequencing and data analysis

Genomic DNA was PCR-amplified using Q5 High-Fidelity 2x Master Mix (NEB M0492) for 20 cycles. For barcoding PCR, 1 µl of the unpurified PCR product was used as a template and amplified for 20 cycles. The barcoded PCR products were further pooled and gel-extracted using Zymo gel extraction kit and DNA clean & concentrator (Zymo research 11-301, 11-303) and quantified by Qubit 1× dsDNA HS assay kits (ThermoFisher Q32851). Sequencing of the pooled amplicons was performed using an Illumina MiniSeq system (300-cycles, FC-420-1004) following the manufacturer’s protocol. The raw MiniSeq output was de-multiplexed using bcl2fastq2 (Illumina, version 2.20.0) with the flag --barcode-mismatches 0. To align the generated fastq files and to quantify editing efficiency, CRISPResso2(45) (version 2.0.40) was used in batch mode with base editor output and the following flags: -w 10, -q 30, -plot_window_size 20, -min_frequency_alleles_around_cut_to_plot 0.1. Indel frequency = (insertions reads + deletions reads)/all aligned reads × 100%. Base editing frequency = A-to-G reads/all aligned reads x 100%.

### Statistical analysis

The statistical analysis was performed in Prism software using either one-way ANOVA with Dunnett correction for multiple comparisons, or two-way ANOVA with Sidak correction for multiple comparisons.

## Results

### Activities and stabilities of previously engineered chemically modified crRNAs

Previously, we developed chemically modified crRNAs that support SpyCas9 activity in cell culture via RNP delivery (23). These crRNAs were heavily or fully modified on the sugar-phosphate backbone with various chemical modifications to enhance stability while retaining significant activity. First, to quickly screen the activity of these crRNAs in a context that mimics viral co-delivery (delivering naked crRNA with virus-expressed effector protein and tracrRNA), we established two stable human HEK293T cell lines that express an effector protein, the tracrRNA and a fluorescent reporter gene (**Figure 1a**) from integrated vectors. In the first case, we generated a clonal line stably expressing SpyCas9 nuclease, tracrRNA, and a traffic light reporter (TLR-MCV1) (40, 46). This reporter consists of a disrupted GFP followed by an out-of-frame mCherry; a double-strand DNA break in the broken GFP region generates a frameshift and places mCherry in-frame, leading to red fluorescence (**Figure 1a, top; Supplemental Figure 1a**). A distinct clonal cell line stably expresses SpyCas9-ABE8e (11), tracrRNA, and a dead_GFP (dGFP) reporter (in which a G-to-A nonsense mutation in the GFP coding sequence prevents GFP expression). Adenine base editing that reverses the point mutation activates GFP expression (**Figure 1a, middle; Supplemental Figure 1b**). We also engineered a third clonal line, derived from mouse Hepa1-6 cells, that stably expresses SpyCas9-ABE8e and tracrRNA (**Figure 1a, bottom**) without a reporter, to facilitate tests at endogenous sites in the mouse genome. In all three cases, the activity of each crRNA chemical modification pattern can be assessed by electroporating the crRNA into the stable cell lines and measuring either reporter activation by Fluorescence Activated Cell Sorting (FACS) analysis, or endogenous target editing by amplicon deep sequencing.

We chose to test a series of the chemical modification patterns of the previously engineered crRNAs that showed robust activity when delivered in the form of the RNP complex (**Figure 1b**) (manuscript in preparation) (47). We synthesized crRNAs with these chemical modification patterns and spacer sequences targeting either the fluorescent reporter gene or endogenous mouse genomic loci. By electroporating the crRNAs into the stable cell lines, we found that the heavily modified C20 showed consistently high activity at all the target sites tested (**Figure 1c-1g**) and significantly improved activity compared to the minimally modified C0 (**Figure 1c**). However, all other heavily and fully modified crRNAs tested showed reduced activity compared with C20, with both SpyCas9 nuclease and SpyCas9-ABE8e, and the activity was inconsistent between different target sites (**Figure 1c-1g**).

Naked crRNA delivered without lipid formulation needs to be stable enough to survive harsh environments *in vivo*. As an initial test of the stabilities of the differently modified crRNAs, we incubated the naked crRNAs shown in **Figure 1b** in non-heat-inactivated fetal bovine serum (FBS) at 37°C. We found that only the fully modified crRNAs (i.e., those without any ribose moieties) resisted ribonuclease degradation and remained stable throughout the 24 hours of incubation (**Figure 1h**). The superior activity but poor stability of C20, together with the opposite trend for C21, C40 and C45, indicate that stability/efficacy trade-offs still exist among these previously engineered crRNAs.

### Protecting oligonucleotides can enhance the potency and stability of heavily modified crRNAs

While continuing the development of crRNA chemical configurations that combine high potency and stability on their own, we explored an alternative strategy to enhance the C20 crRNA’s stability while retaining its consistently high activity across sites (**Figure 1c-1g**). The C20 modification pattern includes six riboses, though each ribose also includes a 3’-phosphorothioate for partial stabilization (**Figure 1b**). To further enhance the stability of C20, we designed a series of fully 2’-*O*-Me modified oligos, which we termed “protecting oligos” (P.O.s), with a sequence complementary to the ribose-containing region of C20. This idea was inspired by the design of DNA/RNA heteroduplex oligonucleotides (HDOs) (48, 49) and by the well-established observation that double-stranded RNAs are more resistant to nuclease degradation than single-stranded RNAs (50–52). Because the P.O. leaves the fully chemically stabilized 3’ portion of the repeat region available for annealing, we hypothesized that it could be displaced by strand invasion of the tracrRNA after its initial annealing to the unprotected crRNA bases, thereby allowing the formation of the active crRNA/tracrRNA complex (**Figure 2a, 2b**) (5).

**Figure 2.**
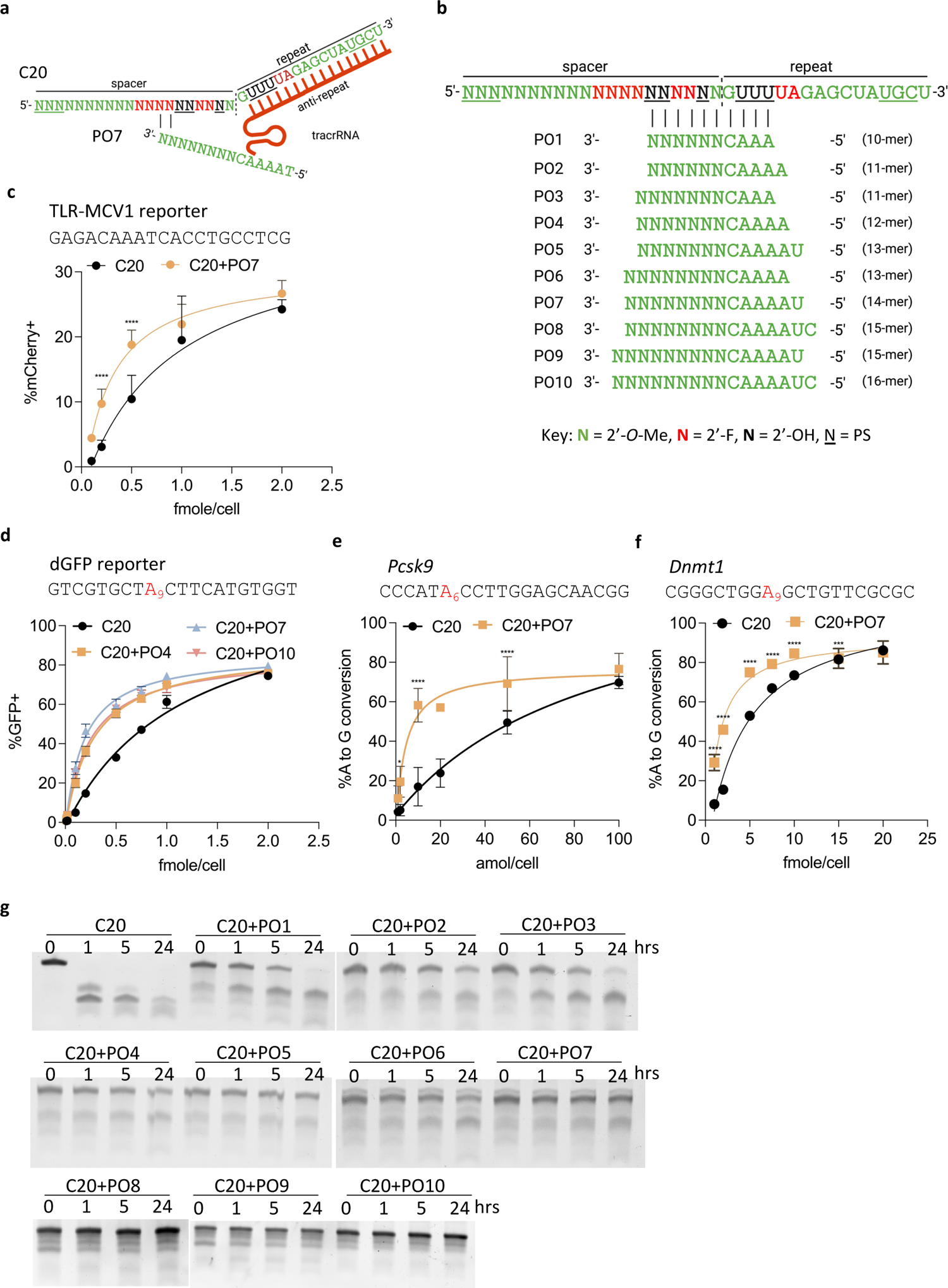
Protecting oligonucleotide (P.O.) enhances the potency and stability of C20 crRNA. a). Schematic representation of the P.O. strategy. Initial annealing of the anti-repeat region of the tracrRNA to the repeat region of the crRNA is hypothesized to lead to P.O. displacement via duplex invasion. b). Schematic representation of the design and modifications of different lengths of P.O.s. c-f). Dose-response curves of C20 targeting different sequences with and without P.O.s in HEK293T-SpyCas9-tracrRNA-TLR-MCV1 reporter (c), HEK293T-SpyCas9-ABE-tracrRNA-dGFP reporter (d), and Hepa1-6-SpyCas9-ABE-tracrRNA stable cell lines (e-f). Editing efficiencies were quantified by FACS (c-d) or targeted amplicon deep sequencing (e-f) (n = 3 biological replicates). Data represent mean ± SD; *, P < 0.05; **, P < 0.01; ***, P < 0.001; ****, P < 0.0001 (two-way ANOVA; C20 was used as control in multiple comparisons). g). The stability of C20 annealed with different lengths of P.O.s (see b) in 10% non-heat-inactivated FBS at 37°C for the indicated numbers of hours. Oligos were resolved by 10% urea-denaturing PAGE followed by SYBR Gold staining.

By annealing the P.O. to C20 and electroporating the duplex into the stable cell lines, we found that the duplexes exhibited significantly enhanced potency compared to C20 alone at all target sites tested (**Figure 2c-2f**). Furthermore, when incubating the duplex with non-heat-inactivated FBS at 37°C, we found that P.O.s greatly improved C20 stability. Furthermore, P.O.s with lengths equal to or longer than 14 nucleotides (nt) can protect the majority of C20 from degradation throughout the 24 hours of incubation (**Figure 2g**). The high efficacy of the C20/P.O. complex further implies that the stability conferred by duplex formation does not prevent productive tracrRNA annealing, as we hypothesized.

We then further explored the mechanisms of P.O. activity. We compared the activities of C20 with P.O.s of different lengths or that anneal to different regions of C20 (**Supplemental Figure 2a**). We found that P.O.s as short as 10 nt (PO1) enhanced C20 potency, although to a lesser extent than the 14 nt P.O. (PO7), possibly because the lower melting temperature limited the degree of C20 protection, as indicated in **Figure 2g**. Conversely, we found that longer P.O.s (up to 22 nt for PO16) reduced C20 potency, possibly due to the strong binding affinity that inhibited initial tracrRNA annealing, P.O. displacement, or both. Furthermore, the positions where the P.O. binds are also important (**Supplemental Figure 2a**). A 16 nt P.O. (B10) that leaves no toe-hold for initial tracrRNA annealing also leads to reduced activity, consistent with our hypothesis that the P.O. needs to be displaced by tracrRNA before C20 can become active (**Supplemental Figure 2a**). Finally, we showed that the P.O. neither inhibited nor enhanced the potency of a fully modified and stabilized crRNA (C40), implying that the potency increase conferred by the P.O. resulted from C20 stabilization (**Supplemental Figure 2b**).

### Protecting oligonucleotides support editing by passive uptake and provide sites for chemical conjugates that enhance *in vivo* distribution

Electroporation bypasses normal steps of cellular uptake and directly introduces the crRNA into the cellular interior (53), enabling functional assessments of editing but providing no information on natural cellular uptake, endosomal escape, or nuclear trafficking. In contrast, it is well established that naked, chemically modified siRNAs and ASOs can be delivered into cultured cells through passive uptake and induce specific mRNA knockdown (18, 54–56). We hypothesized that C20/P.O. complexes can similarly enter cells via passive uptake, enabling endosome escape and nuclear trafficking to form an effector complex and support editing. To test this, we added naked C20, with or without PO7, directly into the cell culture medium of the stable cell line expressing SpyCas9-ABE8e, tracrRNA, and the dGFP reporter (**Figure 3a**). Three days later, we analyzed the editing efficiency by FACS. We found that the unformulated C20/PO7 complex can be delivered via passive uptake in these cells and guide SpyCas9 editing activity (**Figure 3b**).

**Figure 3.**
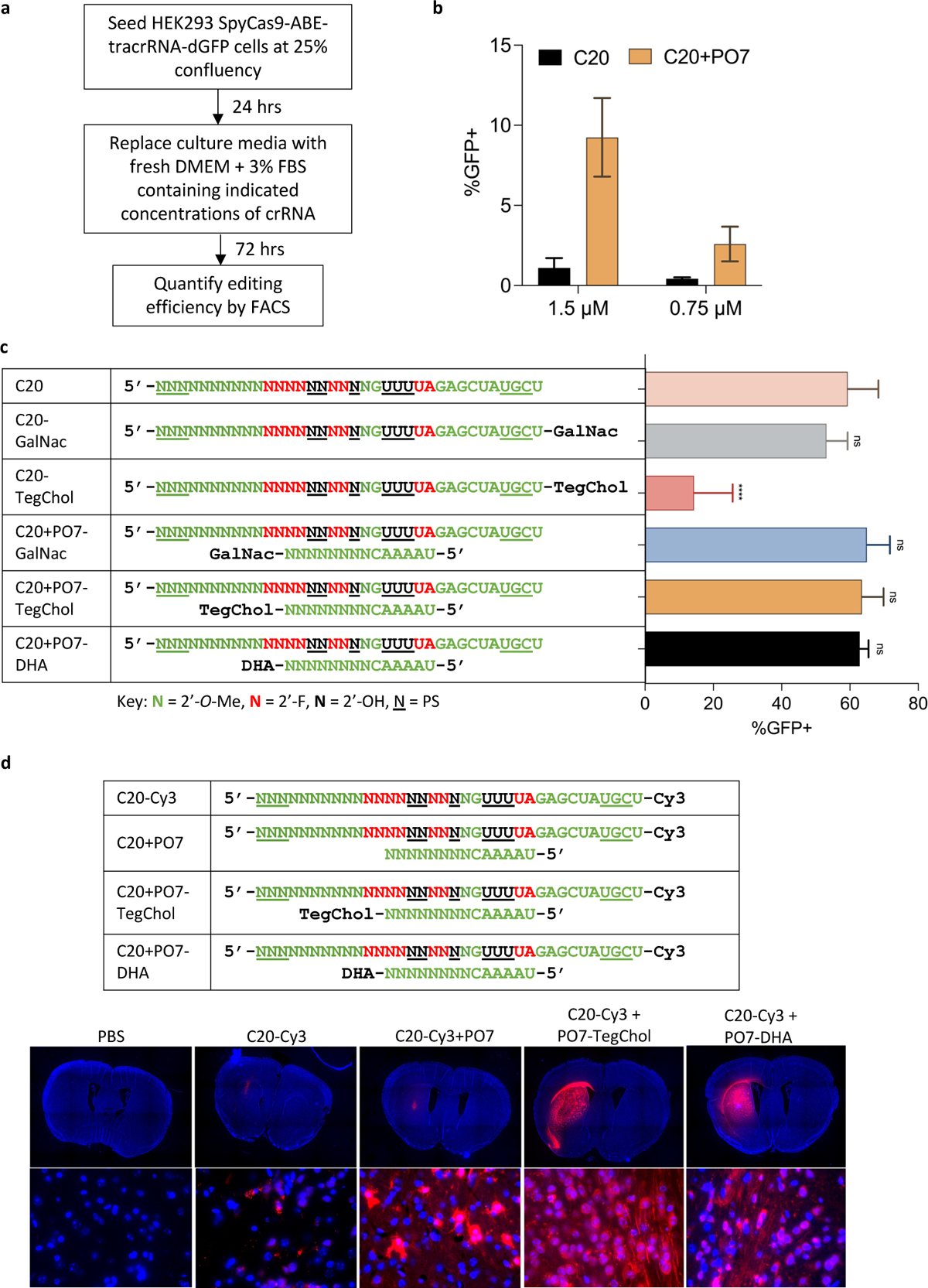
P.O.s promote editing by C20 passive uptake and support a wide range of conjugates for enhanced *in vivo* uptake and distribution in the adult mouse CNS. a). Schematic representation of the experimental process that tests the editing by C20 (with and without P.O.) by passive uptake. b). Editing efficiency by passive uptake (in the absence of transfection or electroporation) of C20, with and without P.O., added to the culture media at the indicated concentrations. Base editing efficiency was measured using the HEK293T-SpyCas9-ABE-tracrRNA-dGFP reporter and FACS analysis. c). 100 pmol C20, with and without conjugates, P.O.s, or both (as shown), were electroporated into 5 x 10^4^ HEK293T-SpyCas9-ABE-tracrRNA-dGFP reporter cells. Editing efficiencies were then measured by FACS analysis. GalNac, trivalent N-acetylgalactosamine; DHA, docosahexaenoic acid; TegChol, tetra-ethylene-glycol-linked cholesterol. Data represent mean ± SD; ns, P > 0.05; ****, P < 0.0001 (one-way ANOVA; C20 was used as control in multiple comparisons). d). Top panel: designs and chemical modification patterns of Cy3-labeled oligos. Bottom panel: *in vivo* distribution of Cy3-labeled C20, with and without P.O.s and lipid conjugates as indicated, in mouse CNS after intrastriatal injection (n = 3 mice per group). Images were acquired using a Leica fluorescent microscope. Upper row of images: 5x magnification tile scan; bottom row: 63x magnification. Blue, DAPI; red, Cy3.

A significant advance in the field of oligonucleotide therapeutics has been the development of oligo bioconjugation (29, 57–61), in which oligos are appended with lipids (60–68), vitamins (69, 70), peptides (71), receptor ligands (58, 59, 72), aptamers (73), or antibodies (74, 75). Depending on the target cell type, these modifications can promote bioavailability, cellular uptake, tissue specificity, and safety of the *in vivo* delivered oligonucleotides through various mechanisms such as 1) penetrating the cell by natural transport mechanisms, 2) specifically binding to a cell surface receptor, or 3) interacting non-specifically with the cell membrane (29). For example, the most clinically advanced bioconjugate, *N*-acetylgalactosamine (GalNac), is a ligand for the asialoglycoprotein receptor (ASGPR), which is highly expressed on the membranes of hepatocytes (58, 59, 72). Conjugating GalNac to therapeutic siRNAs has yielded significant improvements in its pharmacological properties (32). Furthermore, conjugation with lipids such as cholesterol or long-chain fatty acids can broaden therapeutic distribution by facilitating plasma membrane association, binding circulating plasma lipoproteins, promoting uptake by interactions with lipoprotein receptors, or combinations thereof (60). More recently, docosanoic acid (DCA) (63) and docosahexaenoic acid (DHA) (62) have been used to improve biodistribution in extrahepatic tissues such as the muscle and brain.

Despite these advances with ASOs and siRNAs, lipid conjugates such as cholesterol cannot be assumed to be compatible with crRNA and Cas9 activity. Our previous study showed that adding a cholesterol conjugate to crRNA significantly reduced both the *in vitro* and the *in cellulo* activity of the RNP complex (23). The reduced efficiency may be caused by either the strong hydrophobicity or the steric hindrance of the cholesterol conjugate, perhaps affecting guide loading, protein function, or both. A potential advantage of the P.O. approach is that it can provide additional “handles” for conjugates, without the conjugates being present in the final, functional RNP complex. To test this idea, we compared the effects of hydrophobic conjugates on editing efficiency when tethered to C20 or to the P.O.. Through electroporation, we observed that while C20 with cholesterol conjugation showed significantly reduced editing efficiency, cholesterol conjugation to the P.O. did not interfere with editing (**Figure 3c**).

To directly examine the effects of P.O. hydrophobic conjugation on C20 tissue distribution *in vivo*, we delivered Cy3-labeled C20 with and without cholesterol- or DHA-conjugated P.O.s in adult mouse brains by intrastriatal (IS) injection. As expected, we found that compared to unconjugated P.O.s, both cholesterol- and DHA-conjugated P.O.s significantly enhanced the *in vivo* distribution of C20 (**Figure 3d**). These data suggest that the P.O. facilitates various types of conjugation and can be used as a delivery vehicle to enhance the biodistribution of crRNA without interfering with Cas9 activity.

### *In vivo* genome editing in adult mouse CNS by co-delivery of AAV and self-delivering crRNA

Encouraged by these results, we tested whether the unformulated C20/P.O. complex can support genome editing by AAV co-delivery *in vivo*. We chose to test this in a double transgenic reporter mouse model, Cas9/mTmG^+/+^. This mouse model was generated by crossing the homozygous Cas9^+/+^ mice with the homozygous mTmG^+/+^ reporter mice (76). The Cas9^+/+^ mouse model constitutively expresses SpyCas9, and the only component requiring AAV delivery is the tracrRNA. The mTmG^+/+^ reporter consists of *loxP* sites on either side of a membrane-targeted tdTomato (mT) cassette, which constitutively expresses red fluorescence, and a downstream membrane-targeted EGFP (mG) that cannot be expressed until the mT cassette is deleted (**Figure 4a**). First, we injected a self-complementary (sc) AAV9 that expresses tracrRNA into adult Cas9/mTmG^+/+^ mouse brain by IS injection. Two weeks later, we performed IS injection again to deliver a naked C20 that targets the two *loxP* sites to generate a segmental deletion of the mT cassette and allow mG expression. We then detected EGFP expression by performing Immunohistochemistry (IHC) on formalin-fixed mouse brain sections using anti-EGFP antibodies (**Figure 4a**).

**Figure 4.**
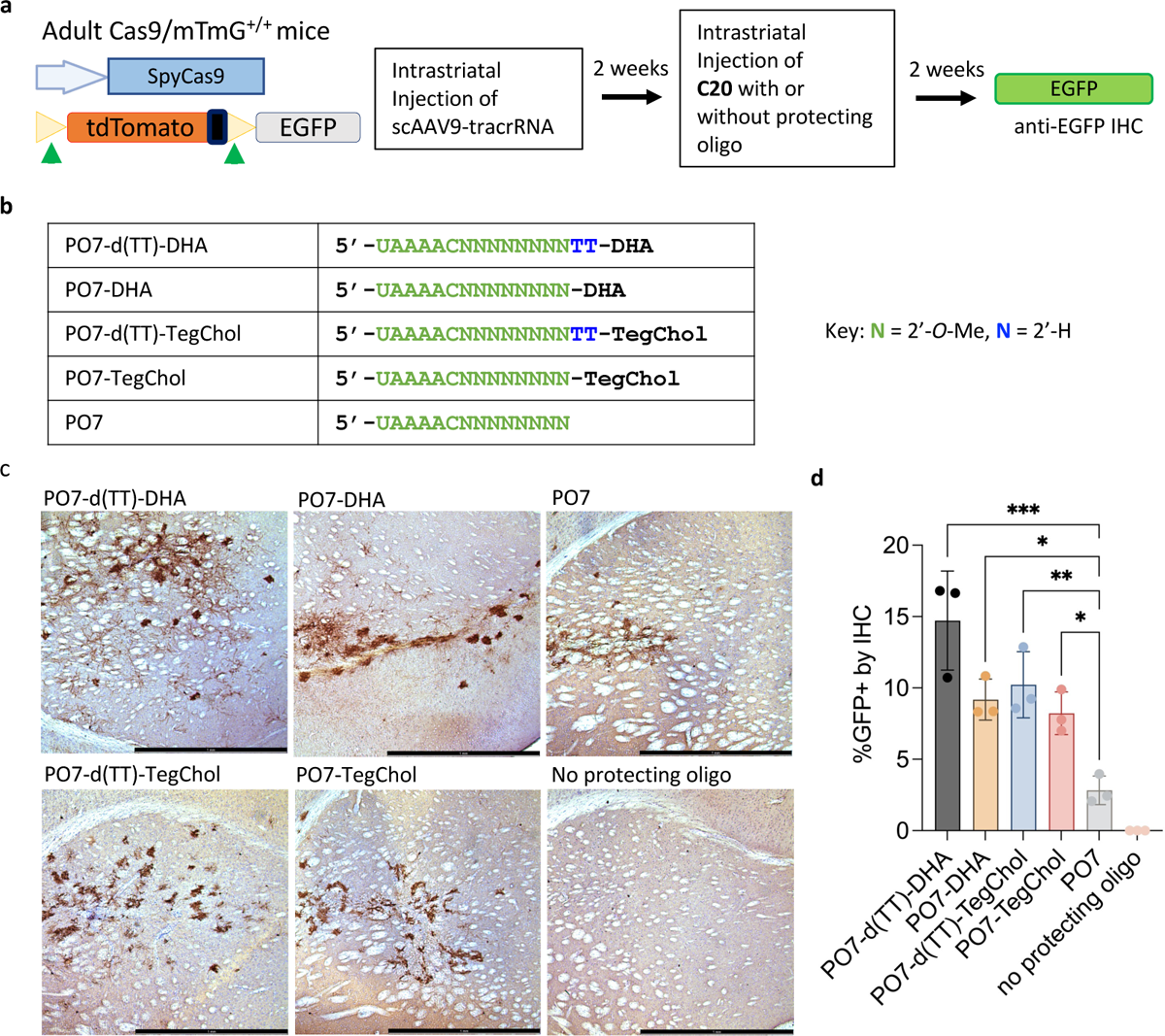
Lipid-conjugated P.O.s promote editing of C20, co-delivered with AAV vector expressing tracrRNA, in the CNS of Cas9-expressing reporter mice. a). Schematic representation of the transgene constructs of Cas9/mTmG^+/+^ double transgenic mouse model and the experimental process to test AAV co-delivery *in vivo* in the mouse CNS. Cleavage at sites flanking the mTmG tdTomato cassette activate EGFP expression. b). Chemical modifications of the P.O.s used in this experiment. d(TT)-DHA, docosahexaenoic acid with a nuclease-cleavable d(TT)-PO-C7 linker; DHA, docosahexaenoic acid; d(TT)-TegChol, tetra-ethylene-glycol-linked cholesterol with a cleavable d(TT)-PO-C7 linker; TegChol, tetra-ethylene-glycol-linked cholesterol. c). Immunohistochemistry (IHC) staining of mouse brain tissue sections using an anti-EGFP antibody. Adult Cas9/mTmG^+/+^ mice were injected with scAAV9-tracrRNA and subsequently injected with C20 and different P.O.s as indicated on the upper left of each image. d). ImageJ quantification of percentages of EGFP positive cells within 1 mm^2^ area around the site of injection (n = 3 mice per group). Data represent mean ± SD; *, P < 0.05; **, P < 0.01; ***, P < 0.001 (one-way ANOVA).

Recent studies on siRNAs showed that compared to cholesterol, conjugation of the less hydrophobic DHA allows broader distribution in the CNS and supports more robust silencing (62). Additionally, using a d(TT)-PO-C7 cleavable linker between siRNA and lipid conjugates has been shown to significantly improve silencing efficacy without altering tissue accumulation (20, 63). This is potentially because the cleavable linker has limited stability *in vivo*, allowing initial tissue distribution while being quickly cleaved after cellular uptake to release the siRNA (preventing oligo attachment to the endosomal membrane or other disruptions to intracellular trafficking). By IHC staining using an anti-EGFP antibody, we found that C20 without P.O. did not generate any editing in the brain (**Figure 4b, 4c**). At the same time, we only observed a few EGFP-positive cells around the injection site with C20 and P.O. with no conjugation. In contrast, C20 with P.O. having either a DHA or cholesterol conjugate (with either cleavable or stable linker) generated more efficient editing with broader distribution (**Figure 4b, 4c**), consistent with the *in vivo* distribution results shown in **Figure 3d**. When quantifying the percentage of EGFP-positive cells within 1 mm^2^ around the site of injection, we found that DHA conjugation performed better than cholesterol conjugation, and having a cleavable linker further improves the editing efficiency (**Figure 4d**).

### *In vivo* genome editing in adult mouse liver by co-delivery of AAV-expressed effector and tracrRNA and GalNac-conjugated, self-delivering crRNA

Next, we tested whether we could achieve *in vivo* genome editing through systemic delivery of C20/P.O. complex co-delivered with AAV-expressed effector protein and the tracrRNA, initially in hepatocytes due to the advanced state of GalNac conjugates in clinical applications of oligonucleotides (58, 59, 72, 77). Here, we chose two established mouse targets to test the efficacy of our co-delivery approach: the *Pcsk9* gene in wild-type B6 mice (**Supplementary Figure 3**), and the *Fah* gene that is mutated in a mouse model of hereditary tyrosinemia type 1 (HT1).

For *Pcsk9* editing in adult B6 mice, we tested C20 with either a 14 nt or a 16 nt P.O. having a trivalent GalNac conjugate (**Figure 5a**) (72). We also included a non-targeting C20 crRNA as well as PBS as negative controls. First, we injected adult mice with AAV to express either SpyCas9-ABE8e and tracrRNA (**Figure 5b**), or SpyCas9 nuclease and tracrRNA (**Figure 5c**) through retro-orbital (RO) injection, and then allowed five weeks for effector protein expression. We then injected C20 with and without GalNac-conjugated P.O. via a single RO injection at 80 mg/kg. The editing efficiency was assessed by extracting genomic DNA from mouse liver and performing targeted amplicon deep sequencing. We found low yet significant editing at the target genomic locus by C20 with both lengths of P.O.s, while no editing was observed for the non-targeting control C20-injected mice (**Figure 5b-5c**).

**Figure 5.**
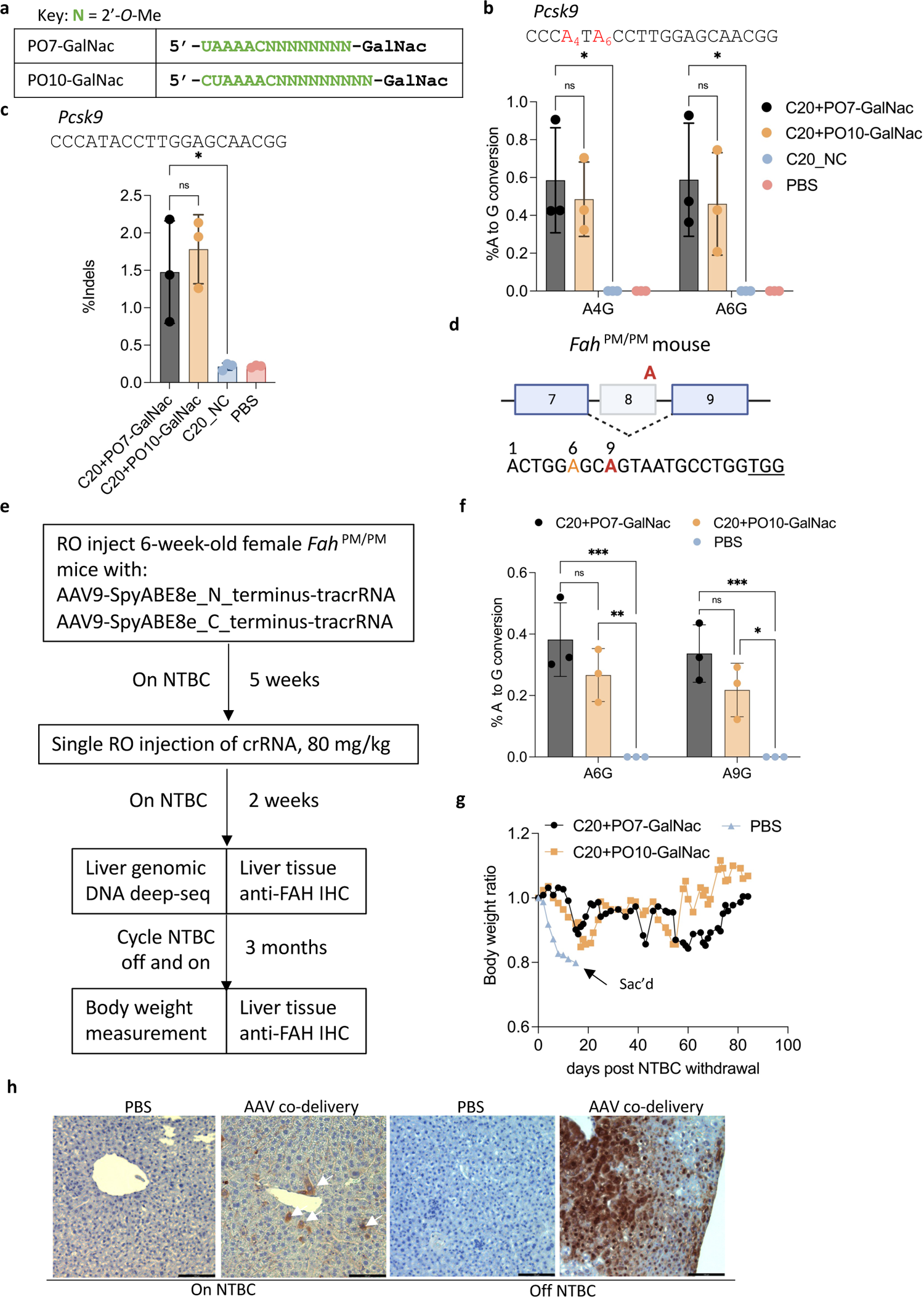
*In vivo* genome editing in the mouse liver following systemic administration of AAV vectors expressing effector + tracrRNA, co-delivered with C20 and GalNac-conjugated P.O.s. a). Chemical modifications of the P.O.s used in this experiment. GalNac, trivalent N-acetylgalactosamine. b). Targeted amplicon deep sequencing quantifying the editing efficiency in the liver by co-delivery of AAV-expressed SpyCas9-ABE8e and tracrRNA, and a single dose of C20 with GalNac-conjugated P.O. at 80 mg/kg. C20_NC, C20 with a non-targeting spacer sequence (n = 3 mice per group). Data represent mean ± SD; ns, P > 0.05; *, P < 0.05, (two-way ANOVA). c). Targeted amplicon deep sequencing quantifying the editing efficiency in the liver by co-delivery of AAV-expressed SpyCas9 nuclease and tracrRNA, and a single dose of 80 mg/kg of C20 with GalNac-conjugated P.O. (n = 3 mice per group). Data present mean ± SD; ns, P > 0.05; *, P < 0.05, (one-way ANOVA). d). Schematic representation of the G-to-A point mutation that generates exon 8 skipping and Fah deficiency in the Fah^PM/PM^ mouse model. The bottom shows a previously validated guide design (44) for SpyCas9-ABE to correct the point mutation. Red, target adenine; orange, bystander adenine; underlined, PAM. e). Schematic representation of the experimental design of *in vivo* AAV co-delivery in the Fah^PM/PM^ mouse model. RO, retro-orbital injection; NTBC, 2-(2-nitro-4-trifluoromethylbenzoyl)-1, 3-cyclohexanedione. f). Base editing efficiency of the genomic DNA from mouse liver treated by AAV-SpyCas9-ABE-tracrRNA co-delivered with the indicated C20/P.O. compound. The editing efficiency was measured by targeted amplicon deep sequencing (n = 3 mice per group). Data represent mean ± SD; *, P < 0.05; **, P < 0.01; ***, P < 0.001 (two-way ANOVA). g). Body weight of the Fah^PM/PM^ mice treated with AAV and the indicated crRNA, measured over 3 months of NTBC off-and-on cycles. h). IHC staining of mouse liver sections using an anti-FAH antibody. White arrows indicate FAH-positive hepatocytes. Scale bar, 100 µm.

Next, we tested whether our co-delivery approach can generate therapeutic-relevant editing by testing in the mouse model of HT1. The HT1 mouse model used in this study is the *Fah*^PM/PM^ mouse, which possesses a G-to-A point mutation in the last nucleotide of exon 8 of the *Fah* gene, which encodes fumarylacetoacetate hydrolase (78, 79). This point mutation generates exon 8 skipping and causes FAH deficiency (**Figure 5d**). Because FAH catalyzes one step of the tyrosine catabolic pathway, FAH deficiency leads to accumulation of toxic fumarylacetoacetate and succinyl acetoacetate, causing damage in multiple organs (78). The *Fah*^PM/PM^ mouse can be treated with 2-(2-nitro-4-trifluoromethylbenzoyl)-1, 3-cyclohexanedione (NTBC), an inhibitor of an enzyme upstream within the tyrosine degradation pathway to prevent toxin accumulation (79). Without such treatment, the mice rapidly lose weight and die. Previously, studies showed that the point mutation in the *Fah*^PM/PM^ mouse could be corrected by SpyCas9-ABE delivered by lipid nanoparticles carrying mRNA and sgRNA, or by plasmid hydrodynamic tail-vein injection (44, 80).

We first injected AAV9 to express SpyCas9-ABE8e and tracrRNA by RO injection, using split inteins to reconstitute the effector as described (81–83). After five weeks, we performed RO injections of C20 with a previously validated spacer sequence targeting the point mutation in the *Fah*^PM/PM^ mouse (44, 80), with either a 14 nt or a 16 nt trivalent GalNac-conjugated P.O., at a single 80 mg/kg dose. The mice were kept on NTBC for two weeks to maintain body weight and allow time for editing to occur. We then sacrificed three mice from each group and extracted liver genomic DNA to measure the editing efficiency by targeted amplicon deep sequencing, as well as detecting FAH-positive hepatocytes via IHC staining using anti-FAH antibodies. For the remaining mouse from each group, we cycled NTBC off and on (7-10 days off followed by 2 days on) for three months to allow FAH-positive hepatocytes to expand, and we monitored body weight to assess functional rescue (**Figure 5e**).

Before NTBC withdrawal, we observed low yet significant on-target editing in the groups that were co-delivered with both AAV9 vectors and the C20/P.O. guides (**Figure 5f**). After NTBC withdrawal, the PBS-injected mouse rapidly lost body weight and was humanely sacrificed after the body weight dropped below 80%. The mice treated with AAV9-expressed SpyCas9-ABE8e and tracrRNA and C20/P.O. duplexes gradually gained body weight over time (**Figure 5g**). The mice were then sacrificed after three months of NTBC cycling, and IHC staining was performed on liver sections using anti-FAH antibodies. We observed the expansion of FAH-positive hepatocytes in the liver tissue treated via AAV9/guide co-delivery (**Figure 5h**). These data provide an initial indication that systemic delivery of C20 and GalNac-conjugated P.O.s, with AAV-expressed effector protein and tracrRNA, can achieve *in vivo* genome editing in the liver and provide therapeutic benefits.

The initial editing efficiency we achieved in the liver, though significant, was low. We then sought to improve the efficiency by optimizing the dosing regimen. The ASGPR, which is expressed on the surface of hepatocytes and is responsible for the uptake of GalNac-conjugated oligos (58, 59, 72), has a cell surface recycling time of 10-15 minutes (77, 84, 85). Data from previous studies also showed that due to ASGPR saturation, hepatocytes can only take up limited amounts of GalNac-conjugated molecules after bolus intravenous injection (86). Our initial study injected a single dose of C20 with GalNac-conjugated P.O. at 80 mg/kg, far exceeding the import capacity of available ASGPR (86). We reasoned that instead of a single large dose, dividing the oligos into three consecutive daily RO injections each with one-third of the previous dose (26 mg/kg) may deliver more oligos into the hepatocyte, thus boosting the editing efficiency.

To test this idea, we first RO-injected AAV9 to express SpyCas9-ABE8e and tracrRNA in the B6 mice. Then, after five weeks, we performed three consecutive daily RO injections of C20 targeting the *Pcsk9* gene complexed with a 14 nt GalNac-conjugated P.O. at 26 mg/kg. We then harvested the mouse liver genomic DNA and performed targeted amplicon deep sequencing to measure the editing efficiency (**Figure 6a**). We found that compared to a single dose of 80 mg/kg oligo, three consecutive daily doses at 26 mg/kg significantly improved the editing efficiency (**Figure 6b**). In addition, we did not observe any sign of toxicity in the liver by redosing, and the oligo-treated mice were in good health, similar to the PBS-injected mice. These data indicate that repeat crRNA dosing is possible and can be well-tolerated, and that further optimization of the dosing regimen promises to provide additional improvements in editing efficiency.

**Figure 6.**
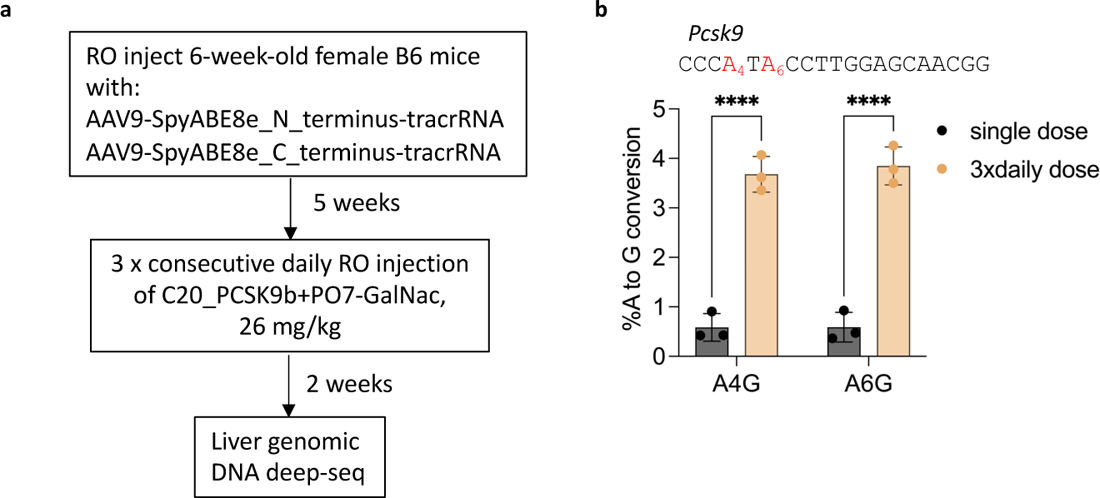
Improving *in vivo* editing efficiency through optimization of the dosing regimen. a). Schematic representation of the experimental process to test a repeat crRNA dosing regimen. RO, retro-orbital injection. b). Comparing the editing efficiency of AAV-SpyCas9-ABE-tracrRNA co-delivered with either a single 80 mg/kg dose, or three consecutive daily injections 26 mg/kg dose of C20+PO07-GalNac targeting the mouse Pcsk9 gene. The editing efficiency was measured by targeted amplicon deep sequencing of mouse liver genomic DNA (n = 3 mice per group). The single-dose data are the same as those shown in Figure 5b. Data represent mean ± SD; ****, P < 0.0001 (two-way ANOVA).

## Discussion

Our results serve as a proof-of-concept study for using P.O. to enhance stability, potency, and cellular uptake of crRNAs to achieve self-delivery *in vivo*. The self-delivering crRNAs can be applied for viral vector co-delivery, potentially allowing multiplexed targeting, especially in tissues that are hard to access (such as the CNS). Tailored co-delivery of the protected crRNA into specific tissues may also provide spatiotemporal control of editing events, reduce off-target effects in unintended organs, and perhaps enable AAV vector inactivation in the future through the co-delivery of vector-targeting guides (**Supplementary Figure 4**) (25).

Although editing efficiencies initially achievable by this approach are still low, our work comprises (to our knowledge) the first description of *in vivo* genome editing via self-delivering crRNA. Importantly, our study also shows that knowledge acquired from the rapidly evolving oligonucleotide therapeutic fields (16, 17, 57, 65, 87) can be readily applied to improve the distribution and efficiency of self-delivering crRNA *in vivo*. Future optimization of P.O.s, including but not limited to additional types of chemical modification, conjugates that allow enhanced or directed cellular uptake and improved pharmacokinetic profiles, and linkers between the P.O. and its conjugates may further enhance potency. Multiple potential directions can be explored that build on these initial proof-of-principle studies. For example, 2’-F residues have been shown to shift the subcellular localization of oligos from the cytoplasm to the nucleus (88), where Cas9 and tracrRNA are localized. Replacing 2’-*O*-Me residues at specific positions within the crRNA or the P.O. with 2’-F may increase effector complex concentration, thus enhancing editing efficiency. Different conjugates can also be applied such as myristic acid (61, 66) and α-tocopherol (69, 70), which were recently shown to provide improved distribution and efficiency properties to ASOs and siRNAs. Additionally, a recent study with siRNAs indicates that a high-affinity universal oligonucleotide anchor with a high-molecular-weight conjugate can reduce the renal clearance of siRNA and enhance blood and tissue retention (89). This approach may also be combined with the P.O. strategy to further enhance potency. Moreover, future studies focusing on optimizing AAV capsids (90, 91), routes of administration (77, 85, 86, 92–96), and dosing regimens for co-delivered AAV vectors and crRNAs promise to further enhance distribution, uptake, safety, and editing efficiency.

Because P.O.s do not interfere with Cas9 activity and tolerate a wide range of bioconjugates, it serves as a delivery vehicle that will be also valuable for delivering fully modified and stabilized crRNA *in vivo*. Furthermore, P.O.s offer opportunities to improve pharmacokinetic and pharmacodynamic behaviors for *in vivo*-delivered, chemically modified crRNA. Our current design of the C20/P.O. complex (a 36 nt crRNA duplexed with a 14 to 16 nt P.O.) mimics asymmetric siRNA designs (a 19 to 21 nt guide strand duplexed with an 11 to 15 nt passenger strand) (16). The single-stranded PS-tails resemble ASO features that serve to enhance cellular uptake. At the same time, the double-stranded region adopts a helical conformation that is relatively consistent across distinct guide sequences, likely facilitating consistency in pharmacokinetic properties. Our P.O. strategy can also be applied to crRNAs with other chemical modification patterns, or to crRNAs for other Cas9 homologs. Finally, the crRNA/P.O. delivery strategy could be extended to transgenic animals or plants that can express the effector and tracrRNA without viral vector co-delivery, advancing genome editing approaches in model organisms and potentially crops or livestock. We anticipate that our study will provide a foundation on which more potent self-delivering crRNA compounds can be engineered for improving the safety and efficacy of *in vivo* CRISPR genome editing via AAV co-delivery.

### Author contributions

E.J.S., H.Z. conceived the study with help from J.K.W. and A.K. H.Z. designed, performed, and analyzed the experiments. K.K. performed mouse injections, with help from H.Z. and N.G. Oligonucleotide synthesis was performed by J.L., D.E., and D.C. R.P. established the HEK293T-SpyCas9-ABE-tracrRNA-dGFP reporter cell line and the Hepa1-6-SpyCas9-ABE-tracrRNA stable cell line. Z.C. purified plasmids for intein-split SpCas9-ABE-tracrRNA AAV packaging. J.X. packaged the AAV9 vectors. G.N. constructed the dGFP reporter. E.J.S., J.K.W., A.K., S.A.W., and D.R.L supervised the research. H.Z. and E.J.S. wrote the manuscript with input and editing from all other authors.

## Data availability

Illumina amplicon deep sequencing data is available at the NCBI Sequencing Read Archive (SRA) database under BioProject ID PRJNA939727. Plasmids first described in this paper will be deposited to Addgene for distribution. All other data are available upon reasonable request.

## Acknowledgments

We thank Nadia Amrani for helpful advice on modified crRNAs, Yueying Cao for *Fah*^PM/PM^ mouse colony maintenance, Julia Alterman for helpful discussion, and Socheata Ly for advice on ImageJ analysis. We also thank the UMCMS Viral Vector Core for AAV packaging services, the UMCMS Morphology Core for tissue sectioning and IHC staining, and the UMCMS Flow Cytometry Core for FACS support. We are grateful to all members of the Watts, Khvorova, Wolfe, Gao, Liu, and Sontheimer labs for their valuable discussions, advice, and helpful feedback.

## Supplementary Figures

**Supplementary Figure 1.**
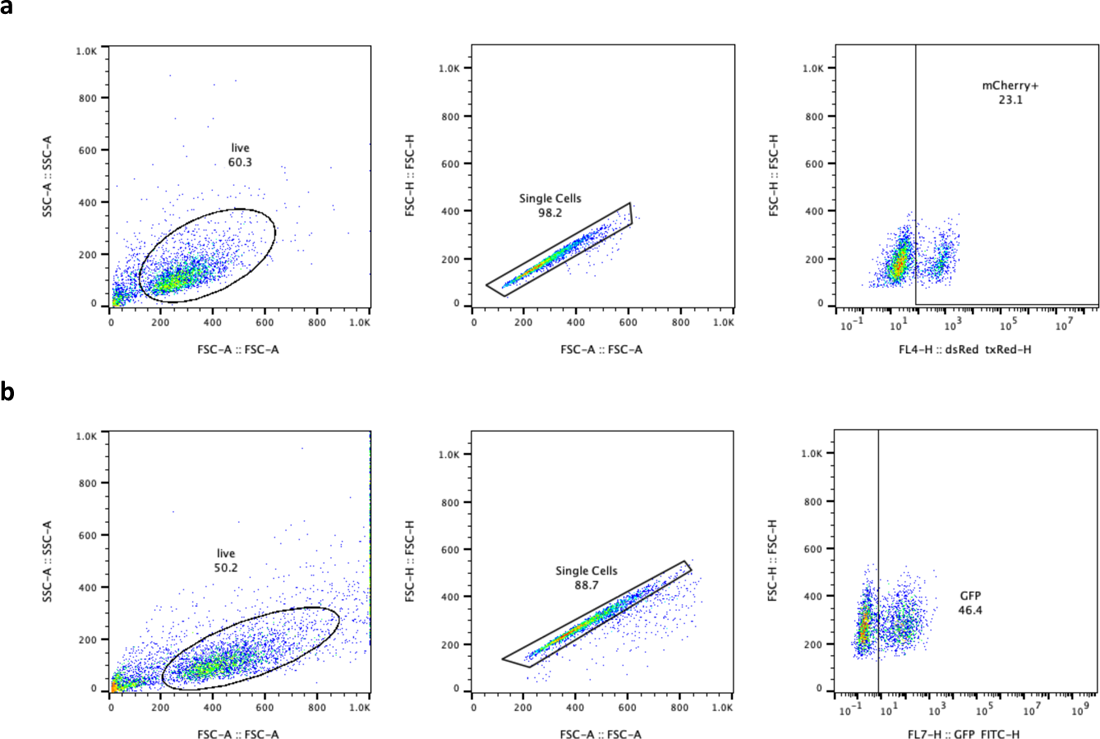
a). Representative flow cytometry gating strategy for the HEK293T-SpyCas9-tracrRNA-TLR-MCV1 stable cell line. b). Representative flow cytometry gating strategy for the HEK293T-SpyCas9-ABE8e-tracrRNA-dGFP stable cell line.

**Supplementary Figure 2.**
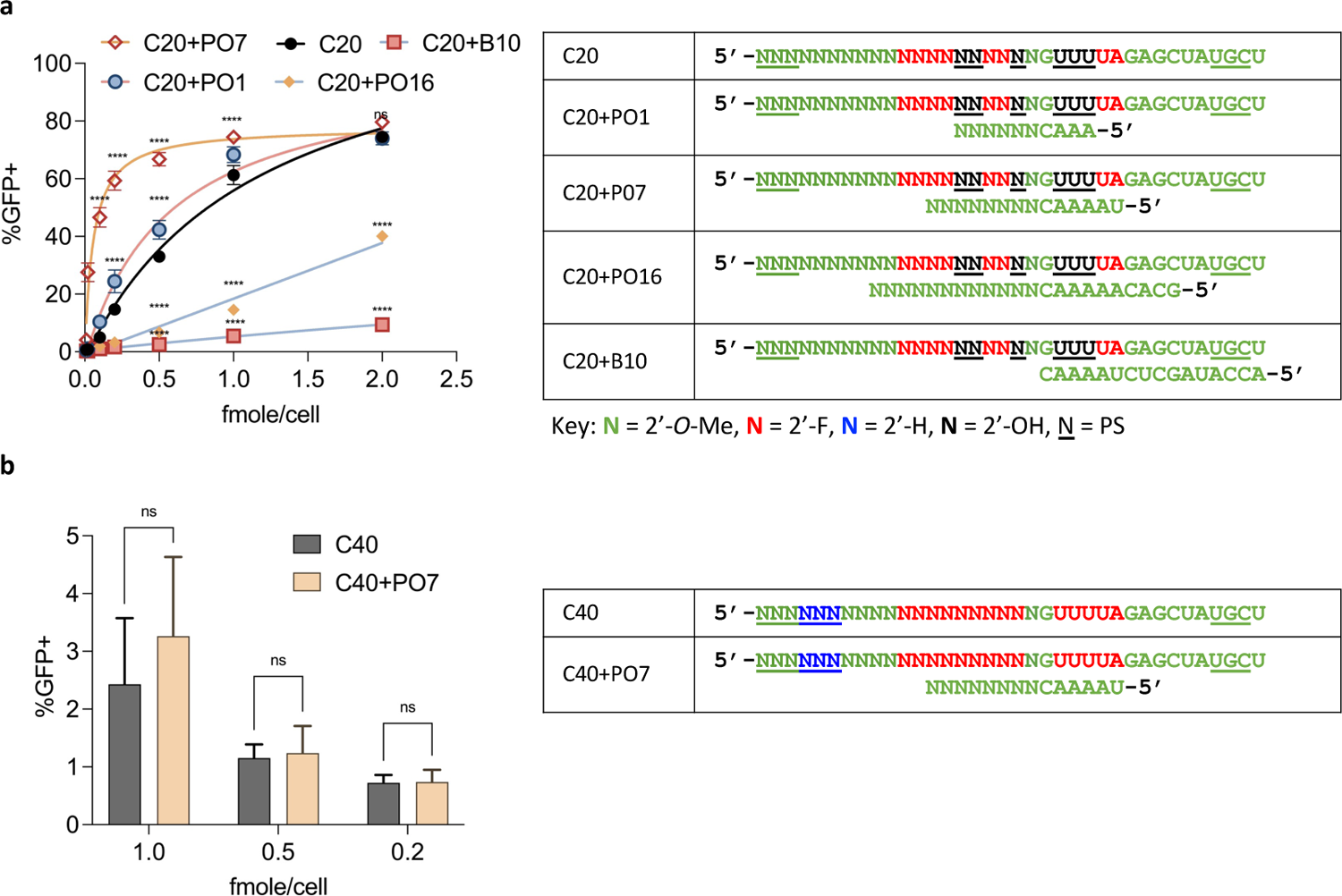
a). Dose-response curves of C20 crRNA, with and without P.O.s, targeting different sequences in the HEK293T-SpyCas9-ABE-tracrRNA-dGFP reporter. Editing efficiencies were quantified by FACS. Data represent mean ± SD; ns, P > 0.05; ****, P < 0.0001 (two-way ANOVA, with C20 used as control in multiple comparisons). The right panel shows the design and chemical modification patterns of tested oligos. b). Editing efficiency of the fully modified crRNA C40, with and without a 14-mer P.O., at different dosages in the SpyCas9-ABE-tracrRNA-dGFP reporter. The right panel shows the chemical modification pattern of the crRNA, and the P.O. Data represent mean ± SD; ns, P > 0.05 (two-way ANOVA).

**Supplementary Figure 3.**
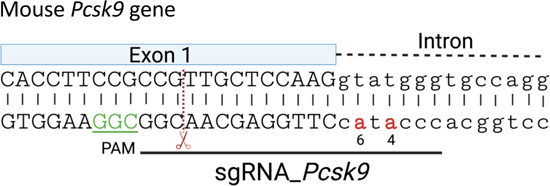
Schematic illustration of the mouse Pcsk9 gene target site used in in vivo co-delivery experiments. The adenines targeted by SpyCas9-ABE8e are labeled in red and bold. The SpyCas9 cleavage site is marked with a red dashed line. The PAM region is underlined in green.

**Supplementary Figure 4.**
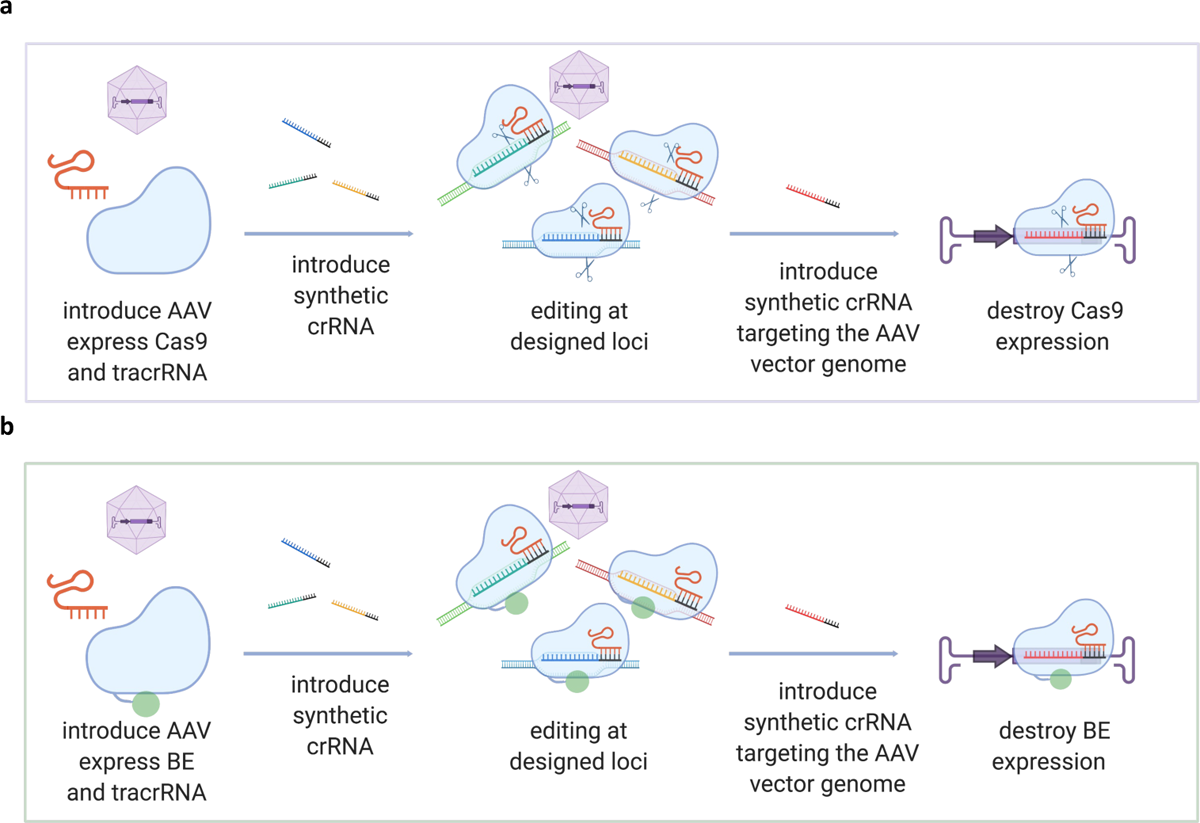
Schematic representation of potential applications of the AAV co-delivery strategy using self-delivering, chemically stabilized crRNA for (a) Cas9 nuclease-based editing or (b) base editing (BE).

## Supplementary Notes

Includes nucleotide sequences of the plasmids first described in this manuscript.

### Lentivector for SpyCas9-tracrRNA transgene integration

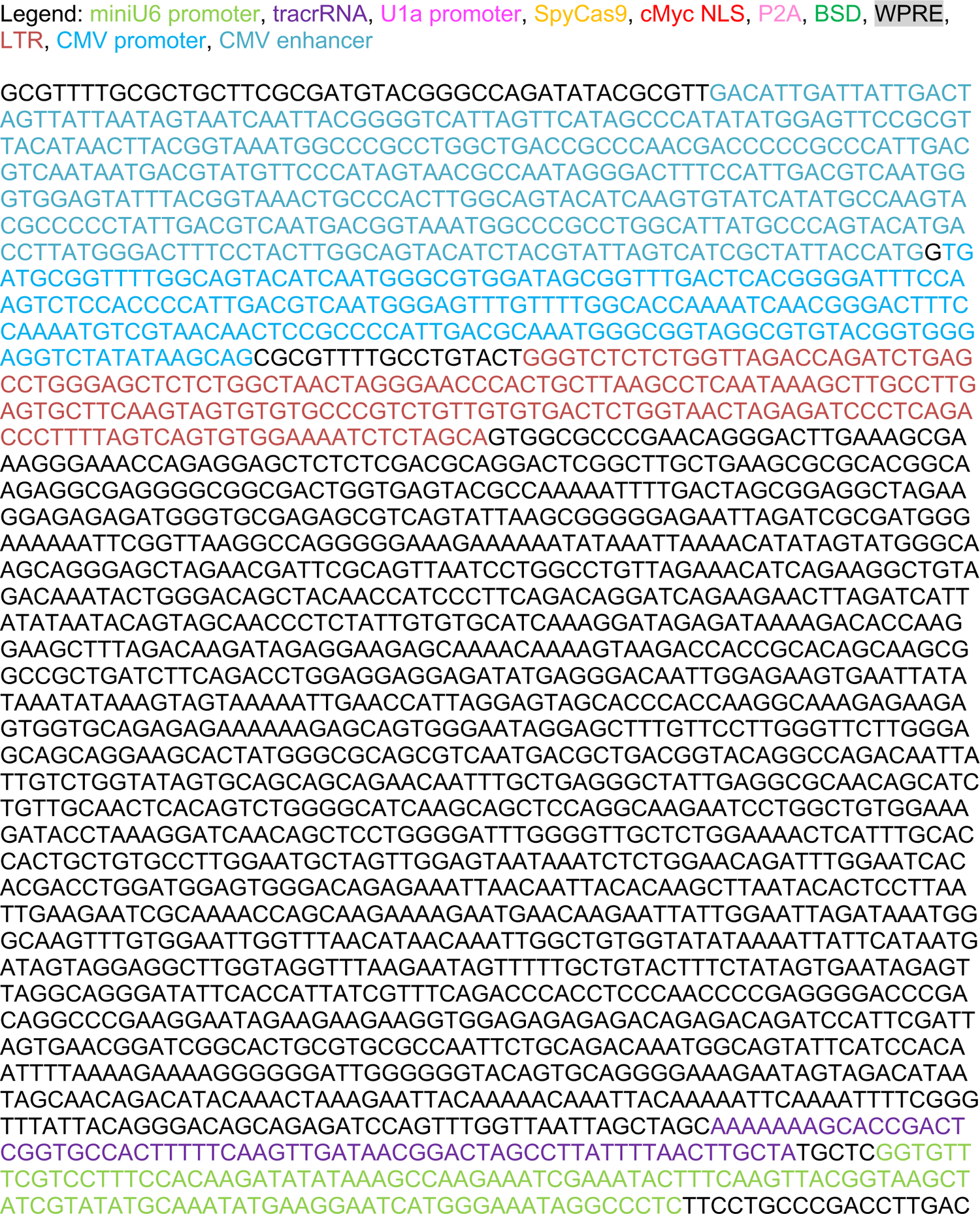
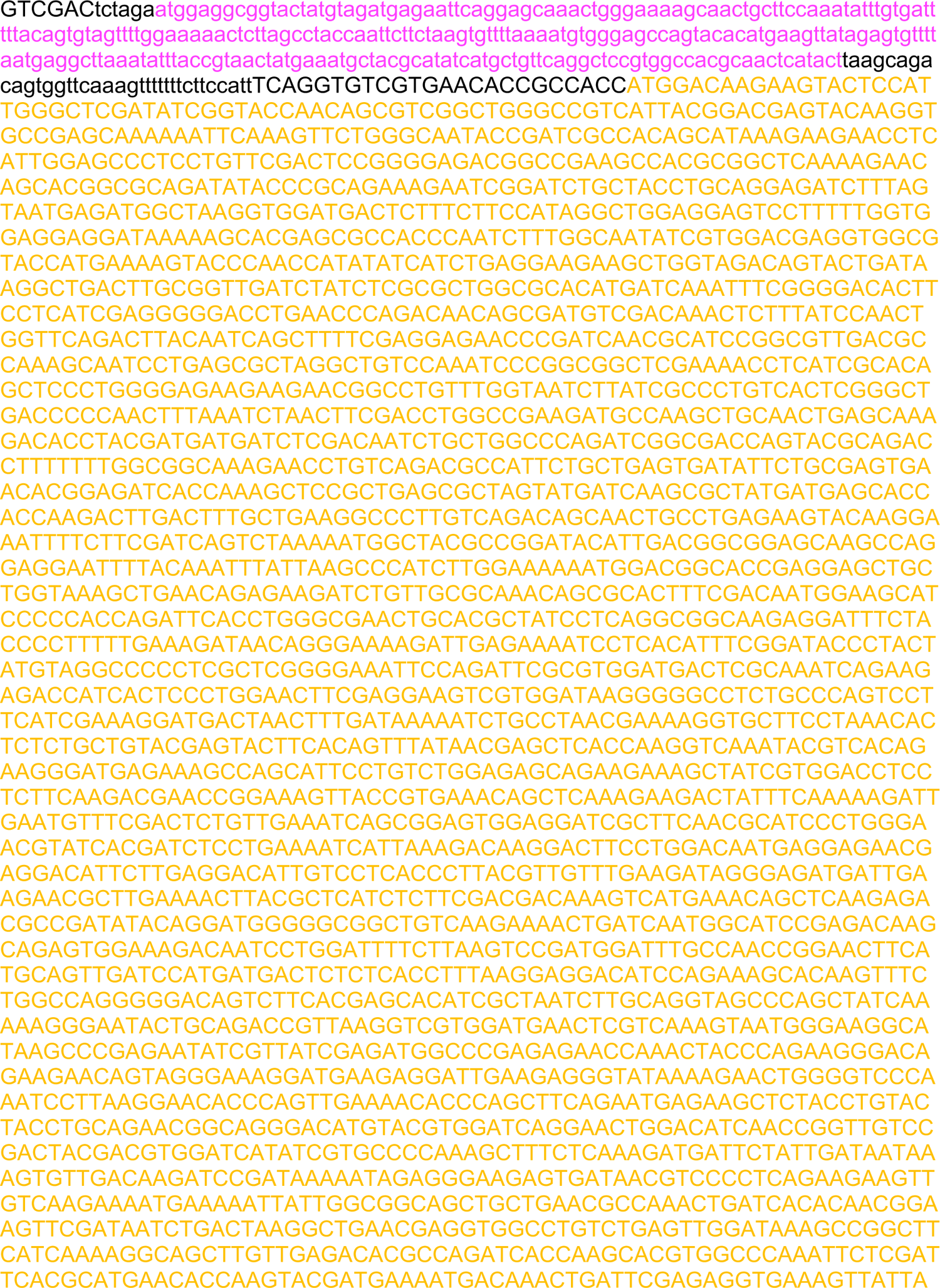
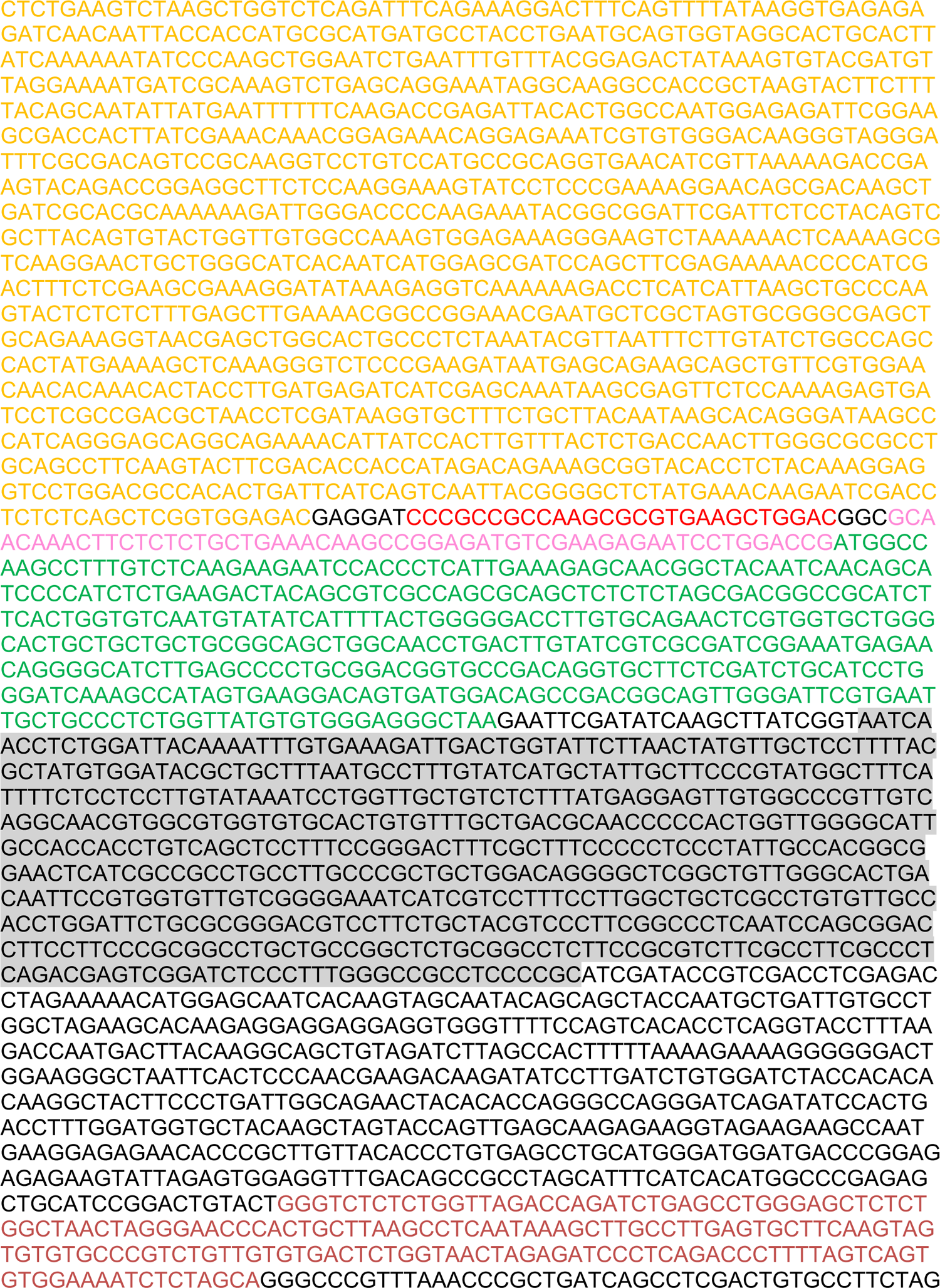
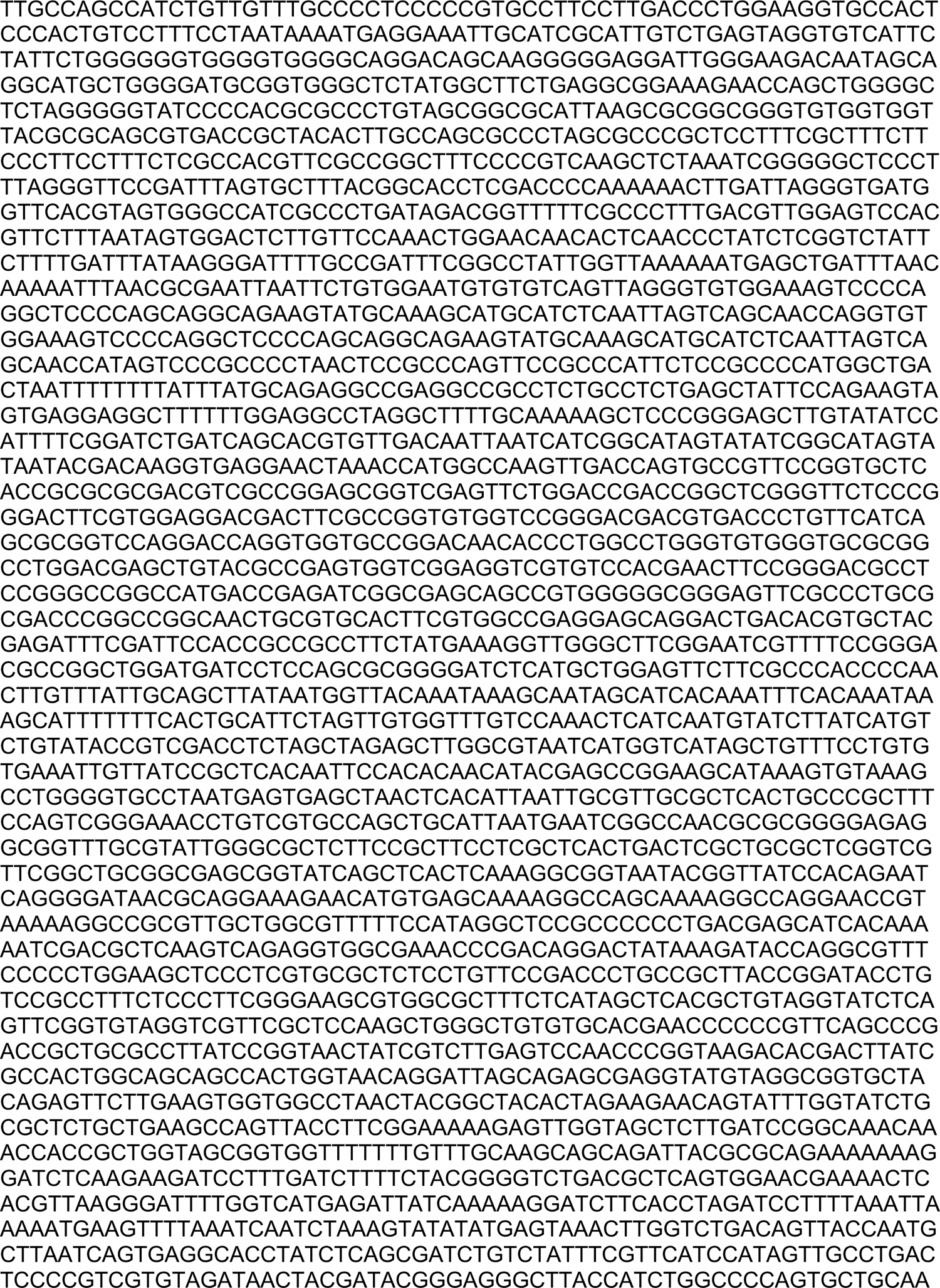
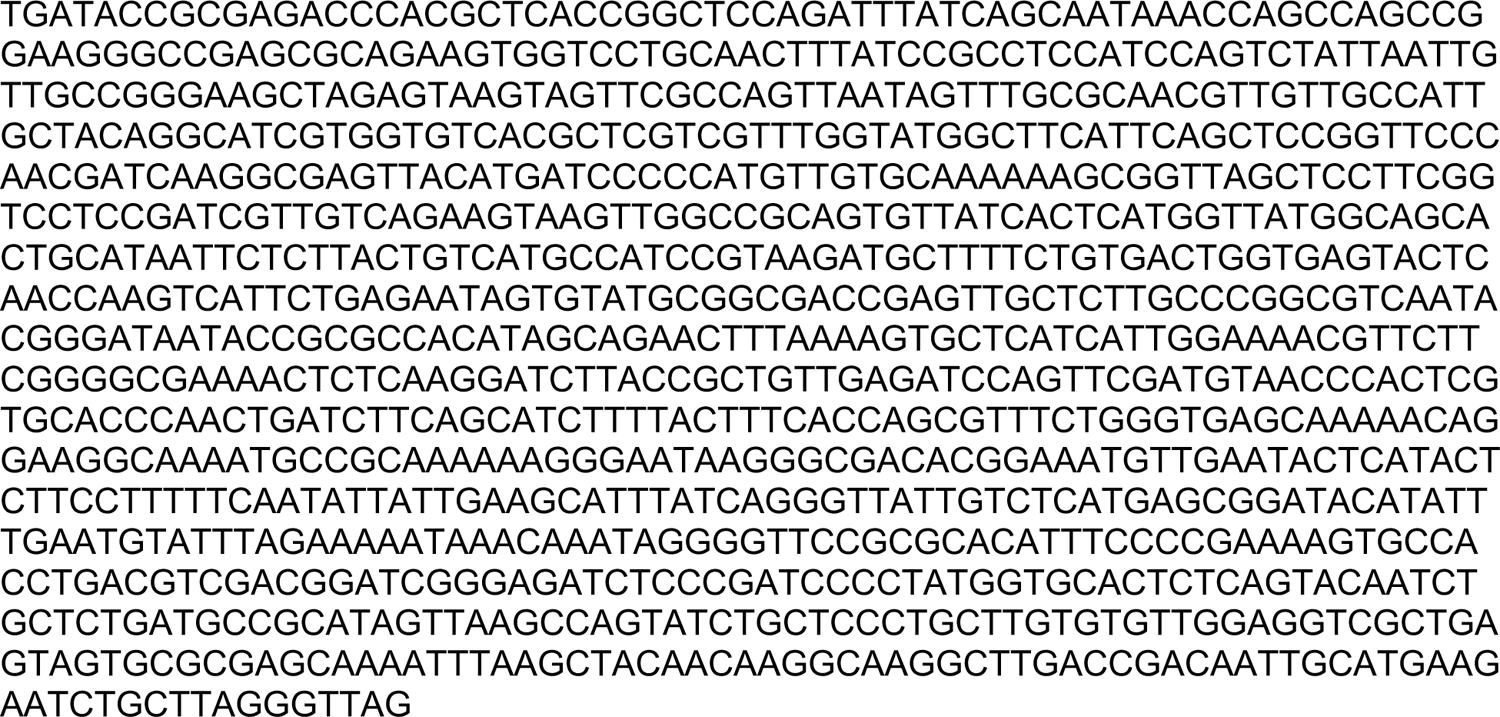

### Lentivector for dGFP reporter transgene integration

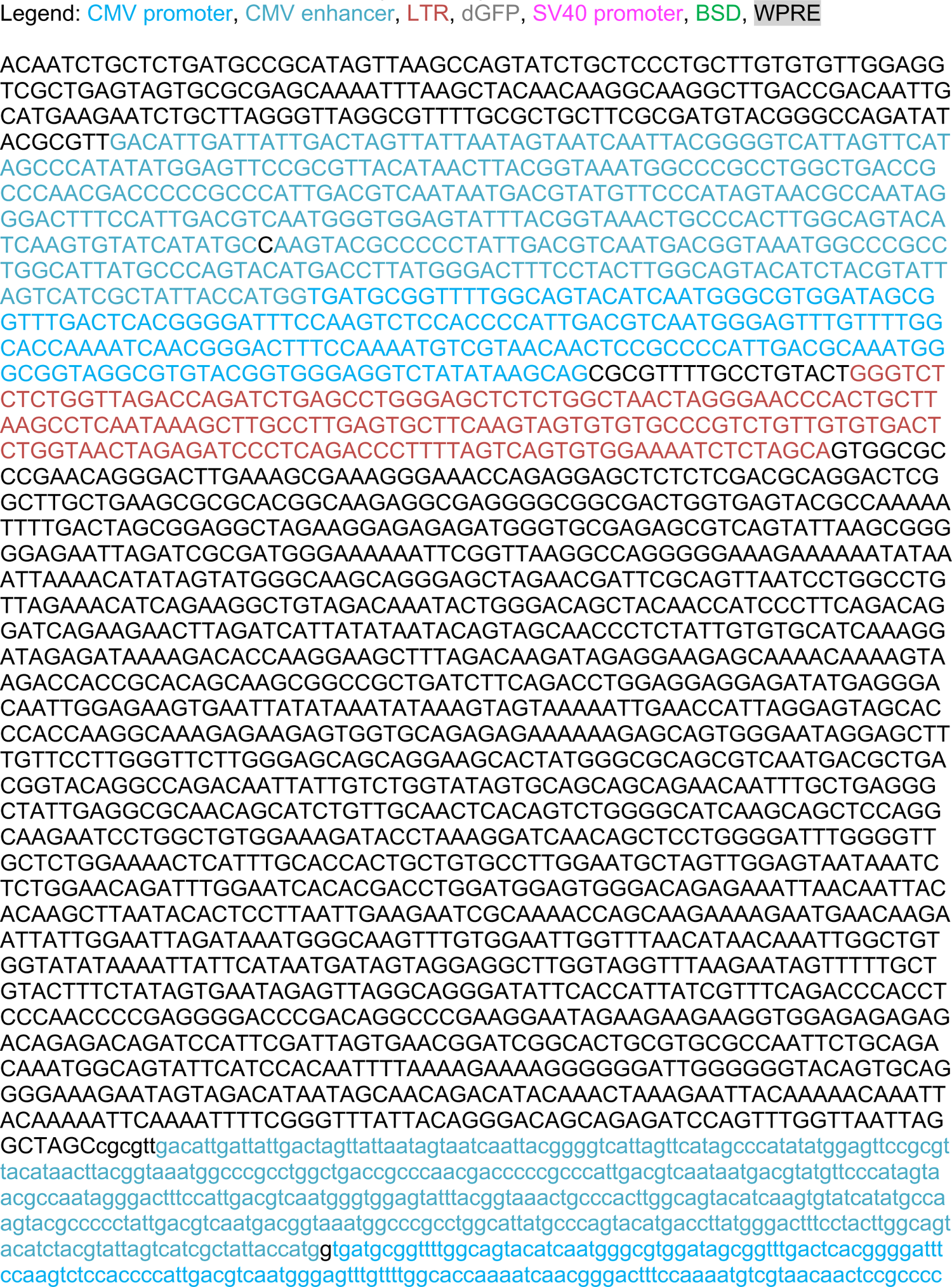
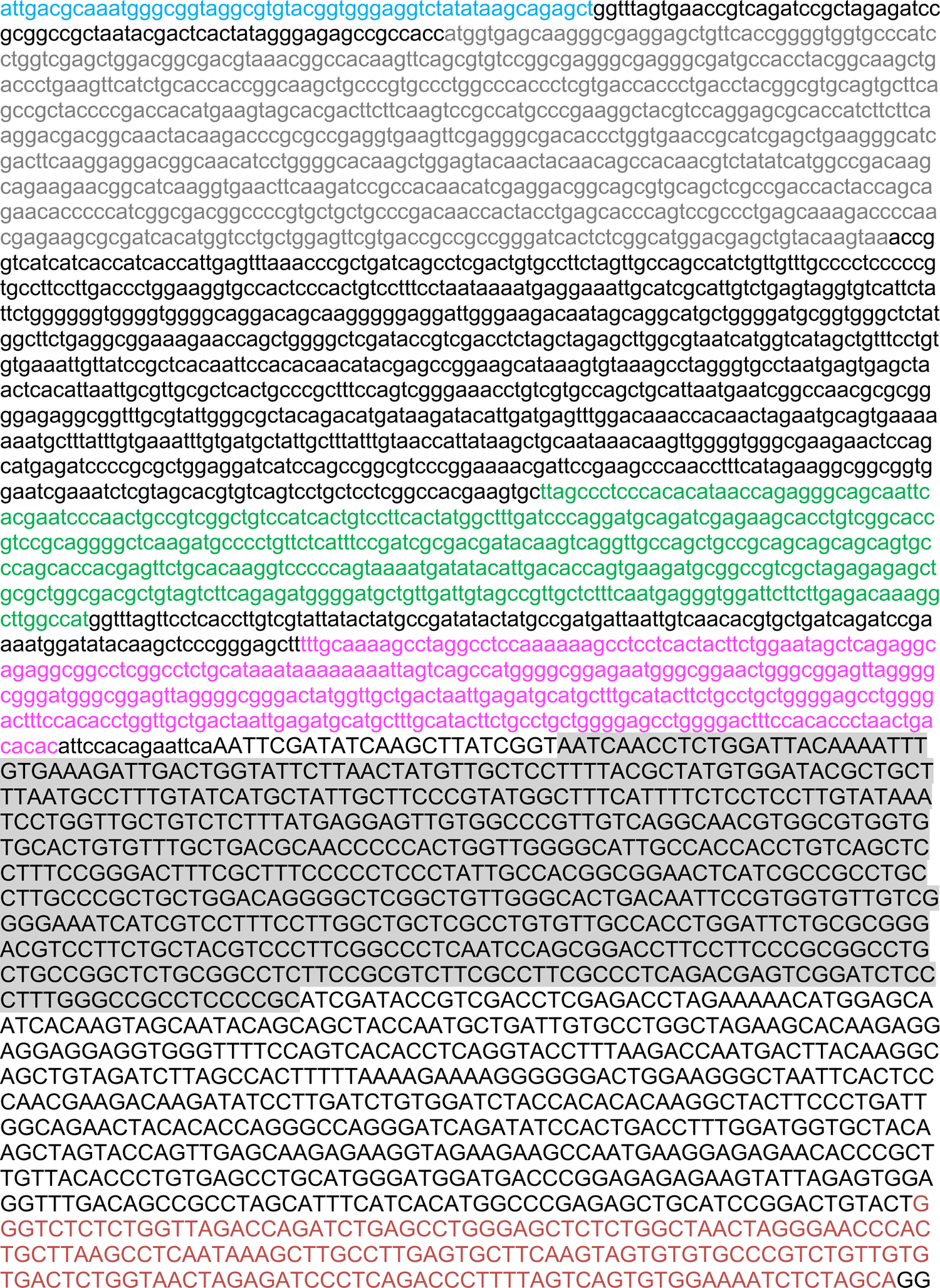
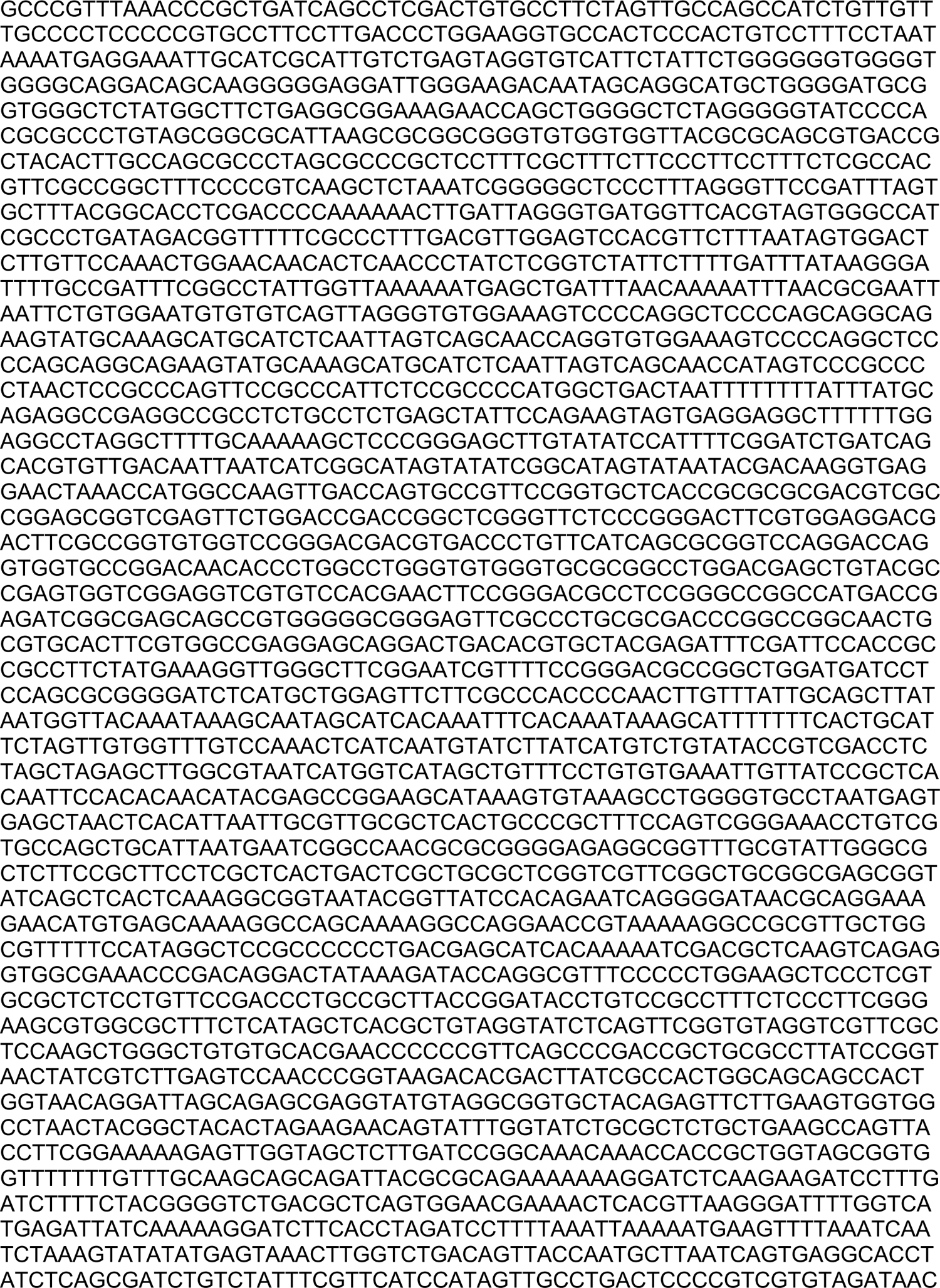
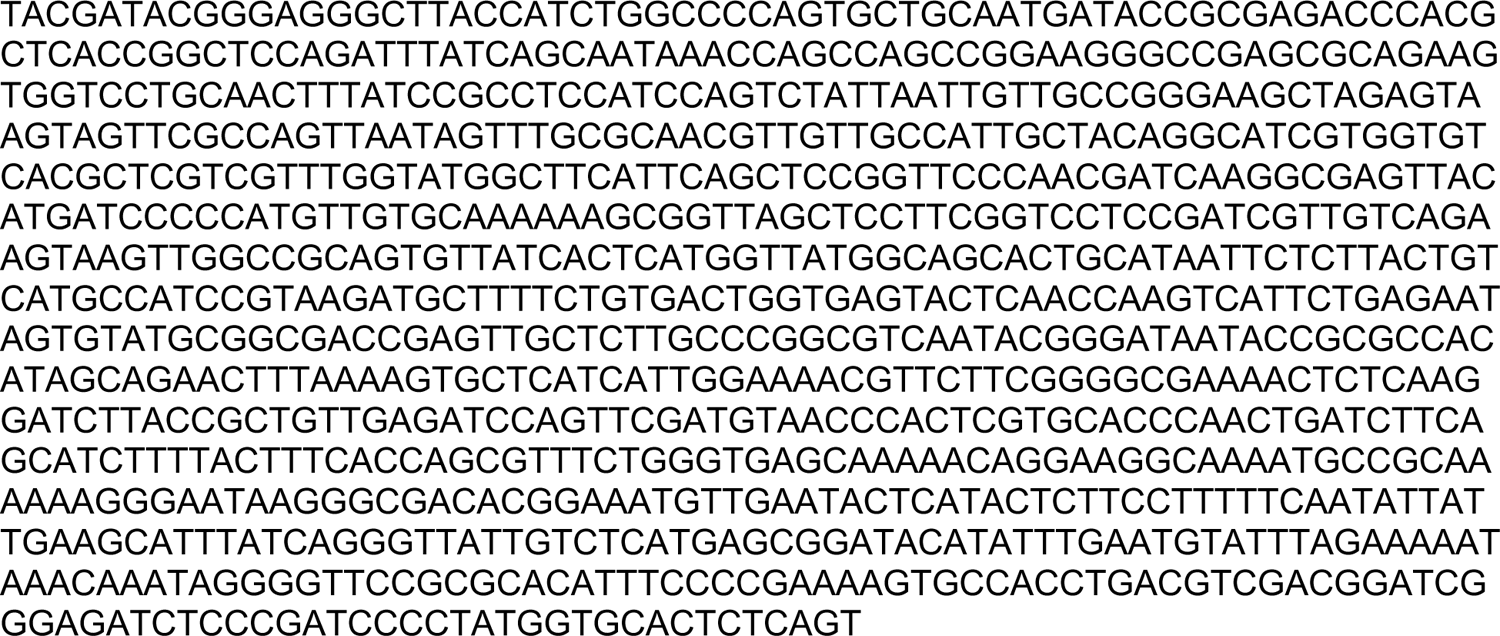

### PiggyBac vector for integration of SpyCas9-ABE8e-tracrRNA

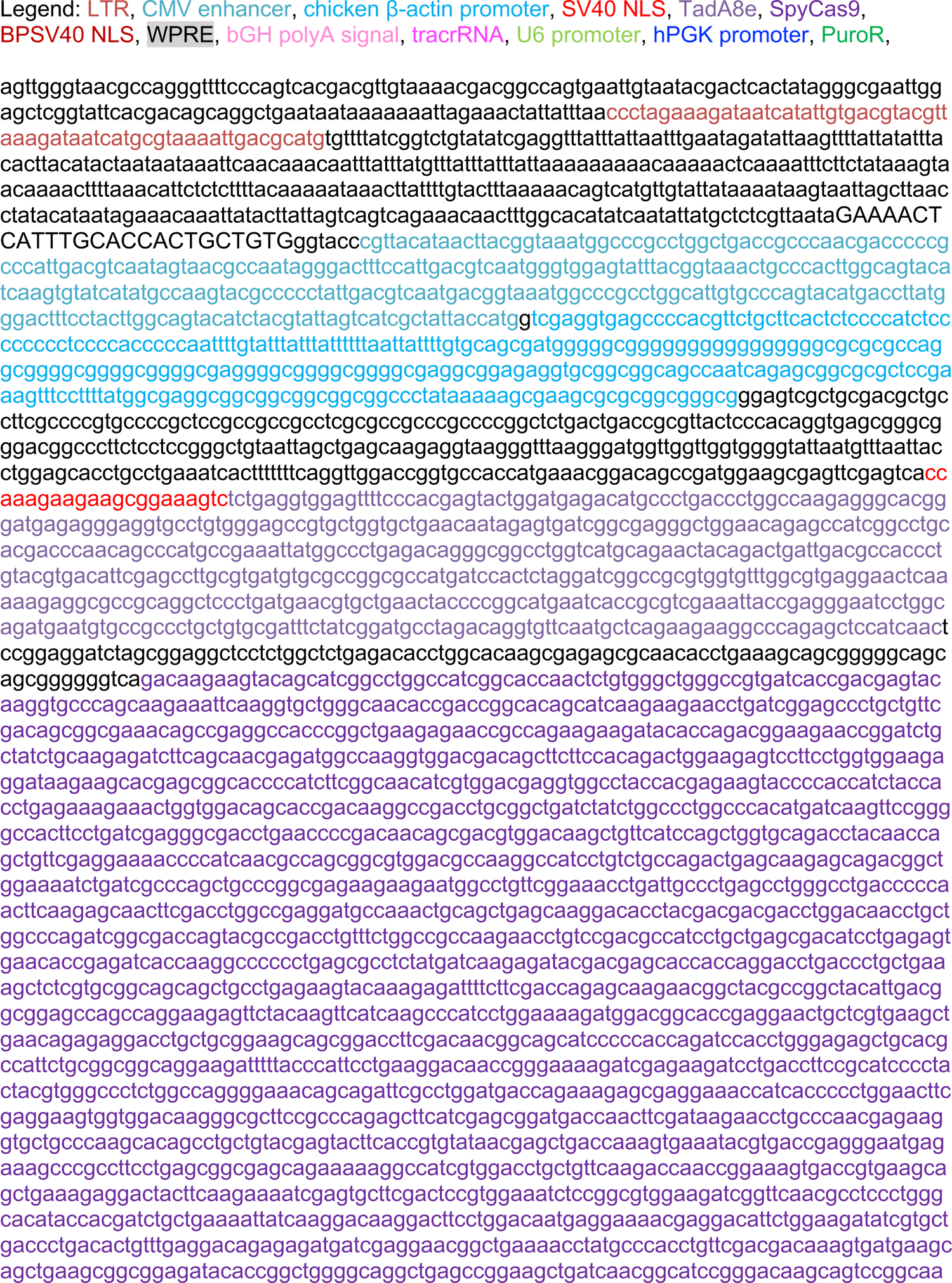
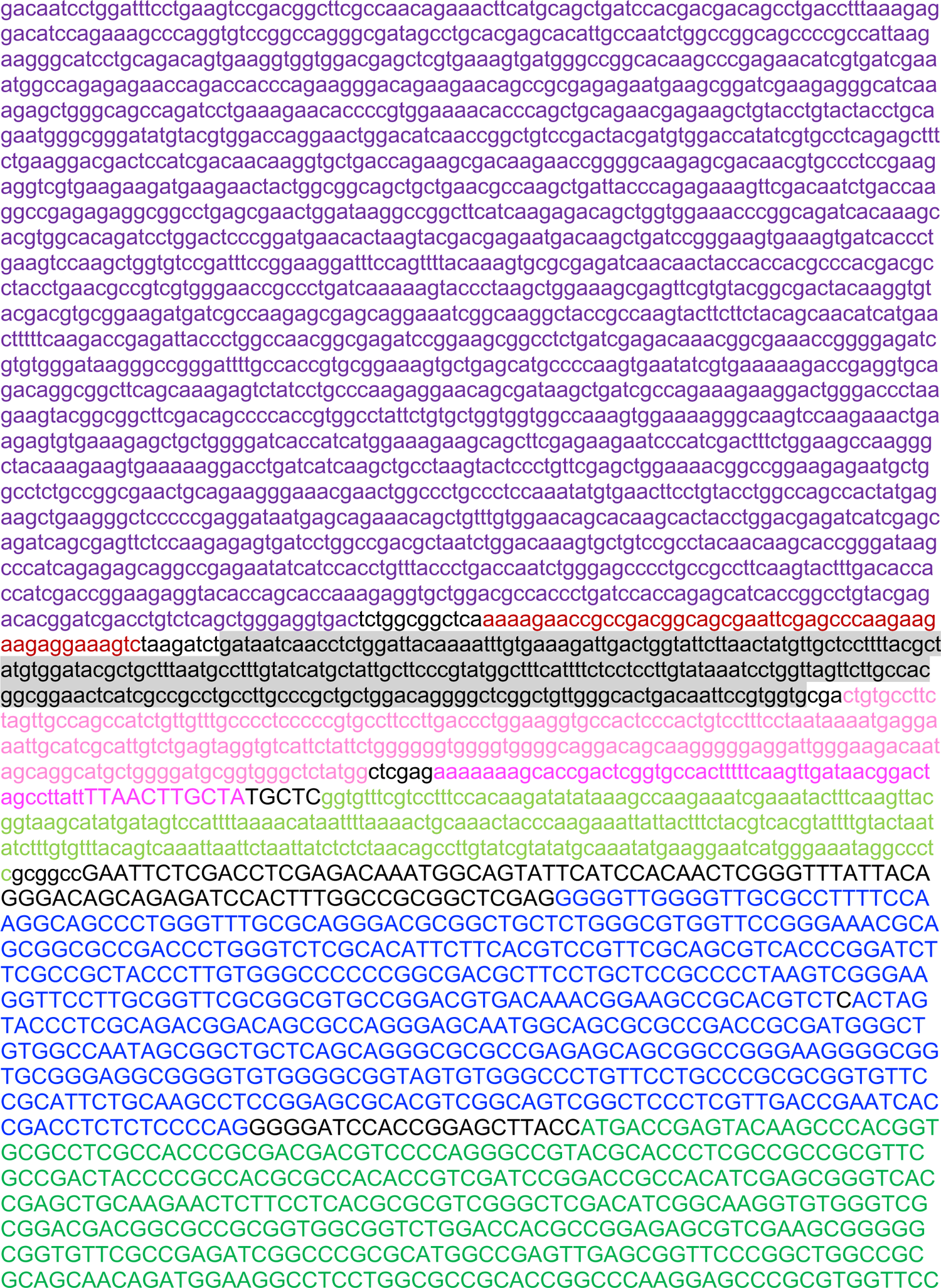
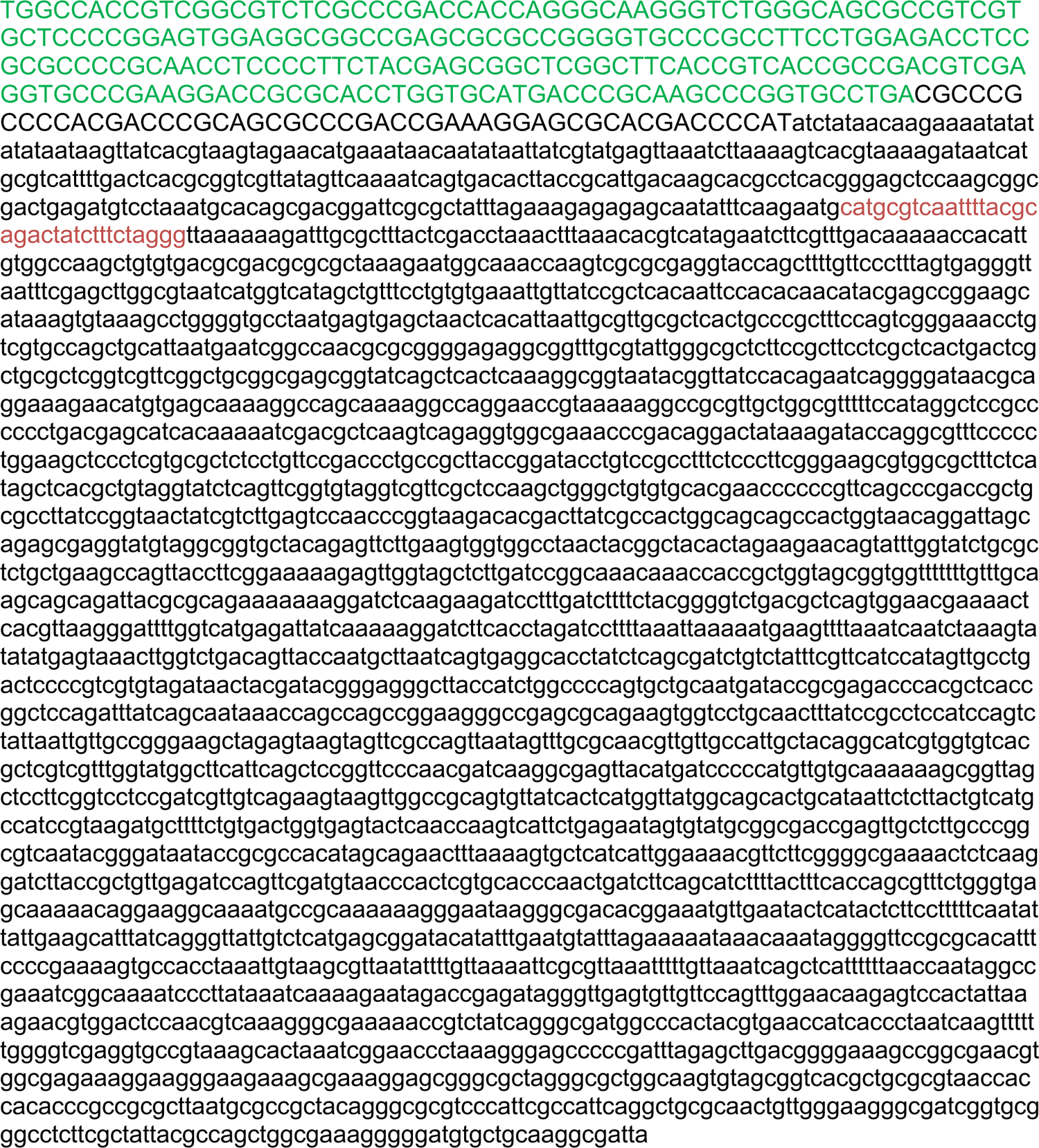

### scAAV-tracrRNA used in Figure 4 and 5c

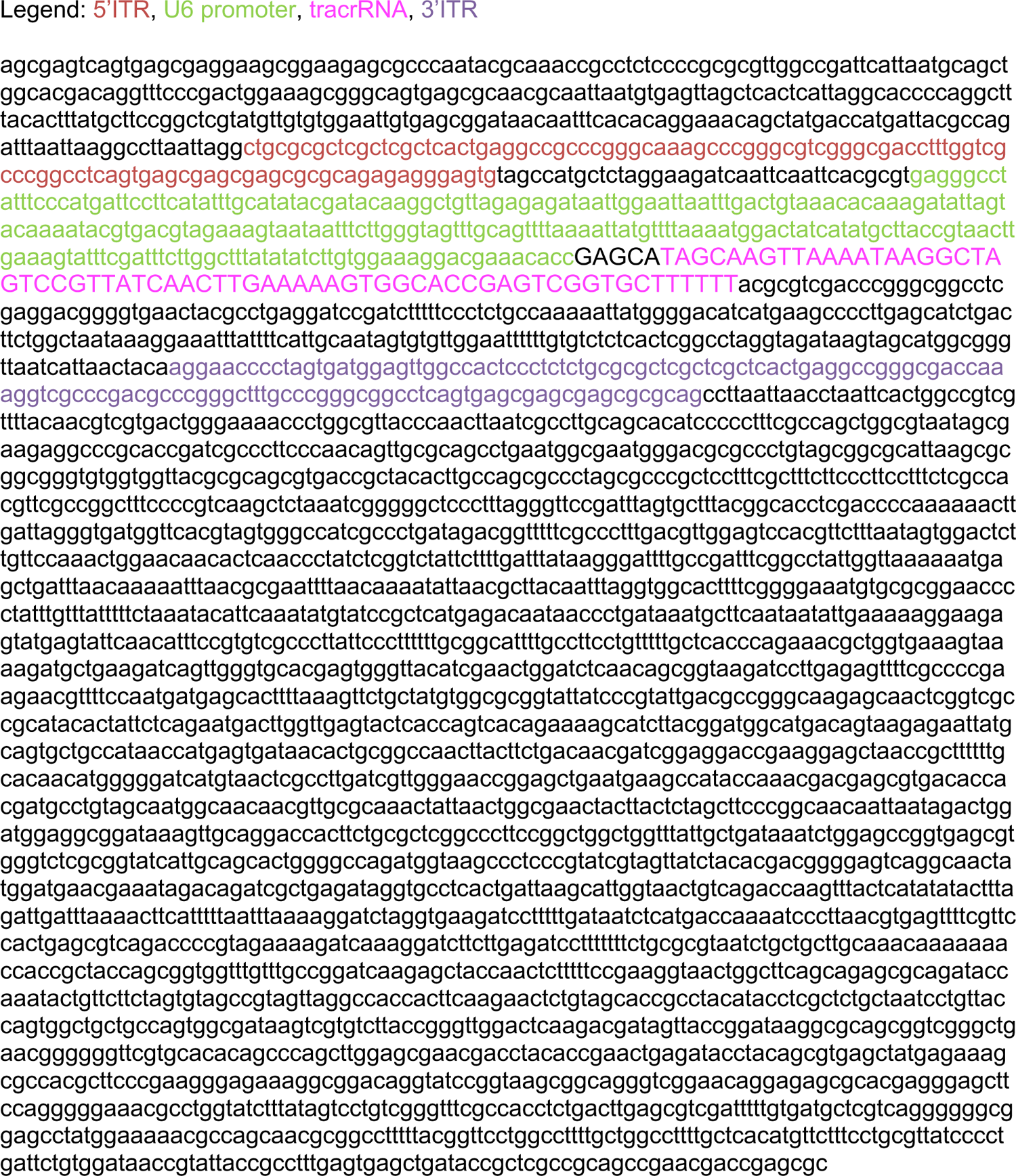

### AAV-SpyCas9-ABE8e-N-tracrRNA used in Figure 5b

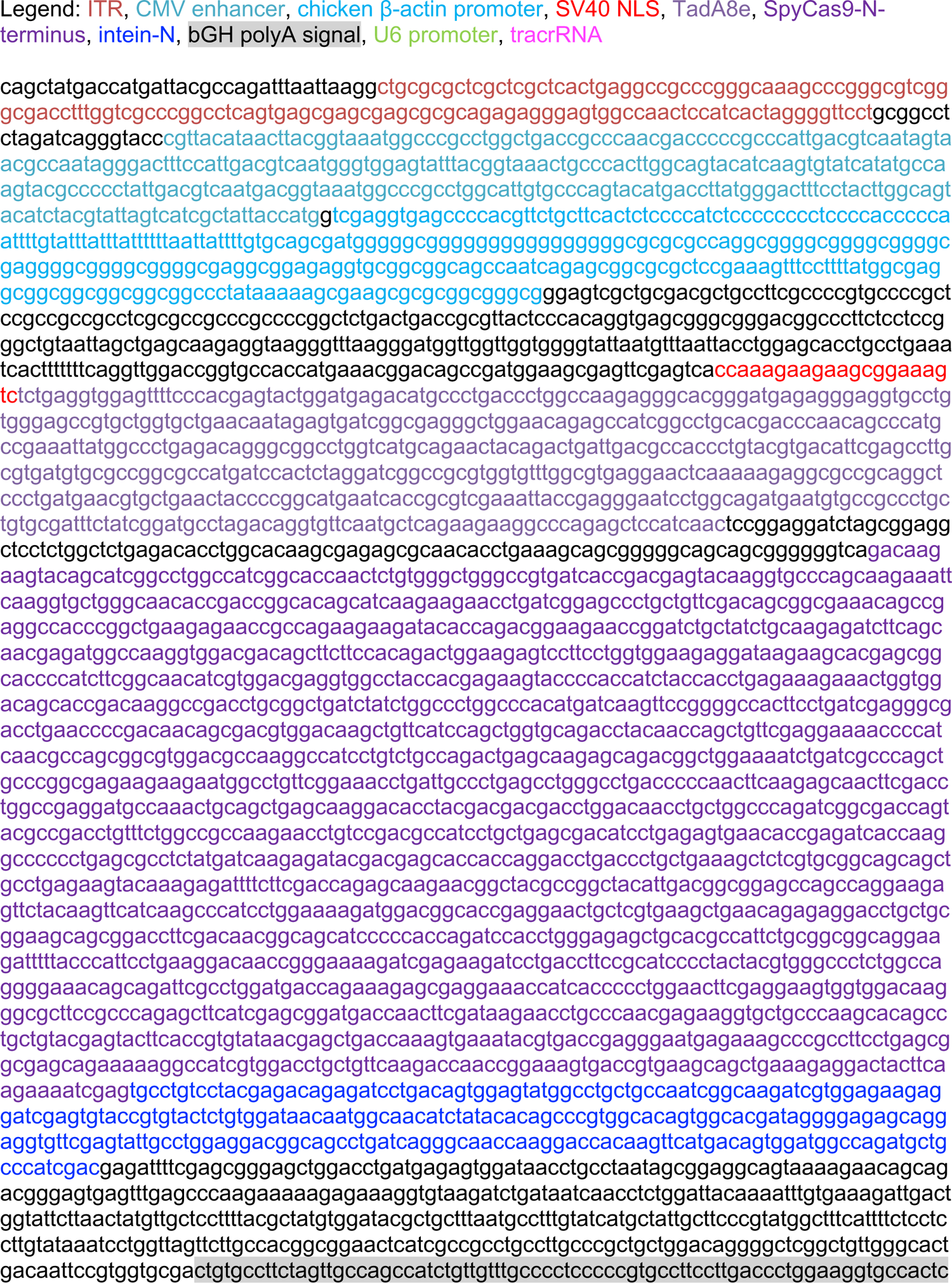
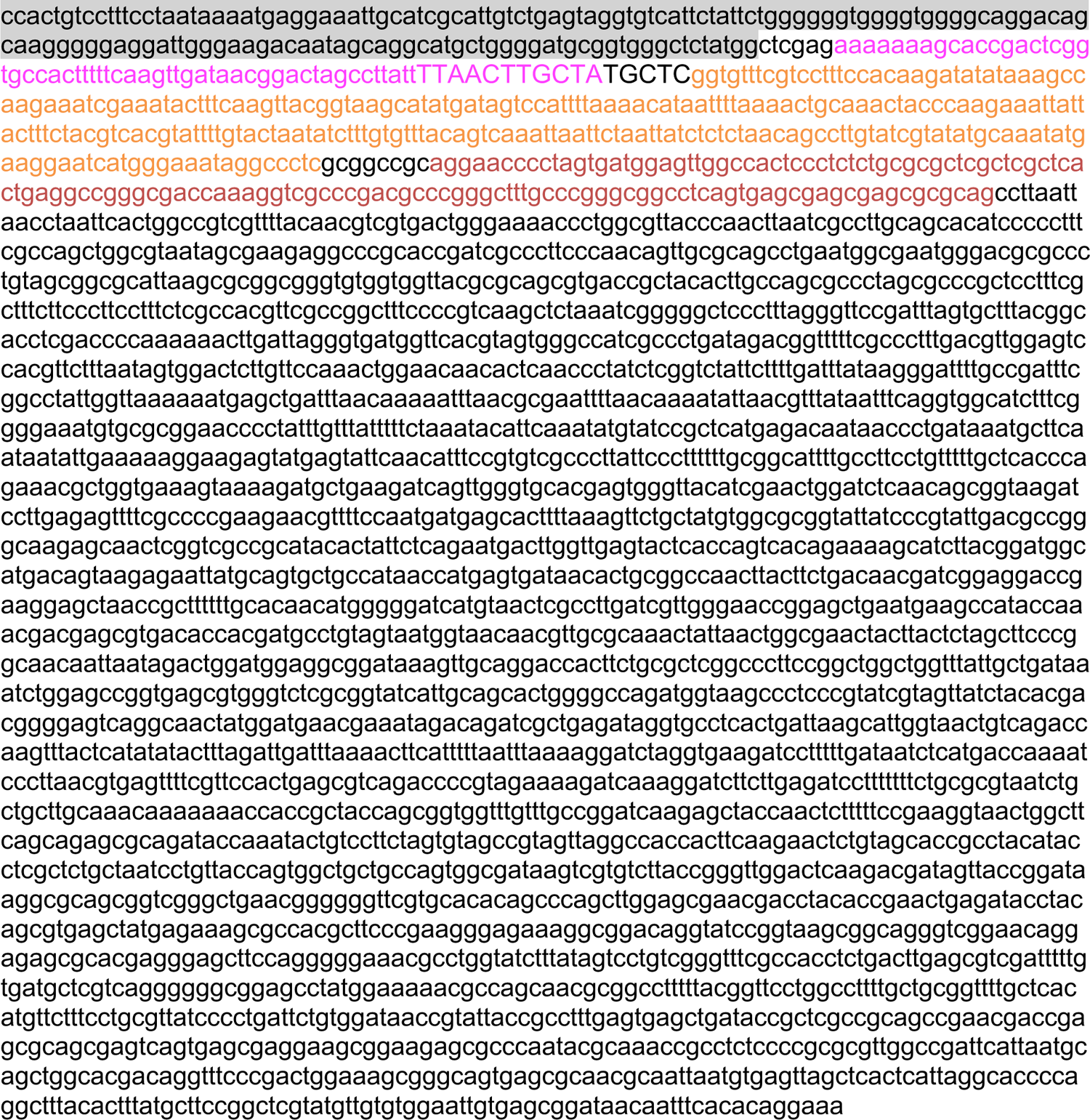

### AAV-SpyCas9-ABE8e-C-tracrRNA used in Figure 5b

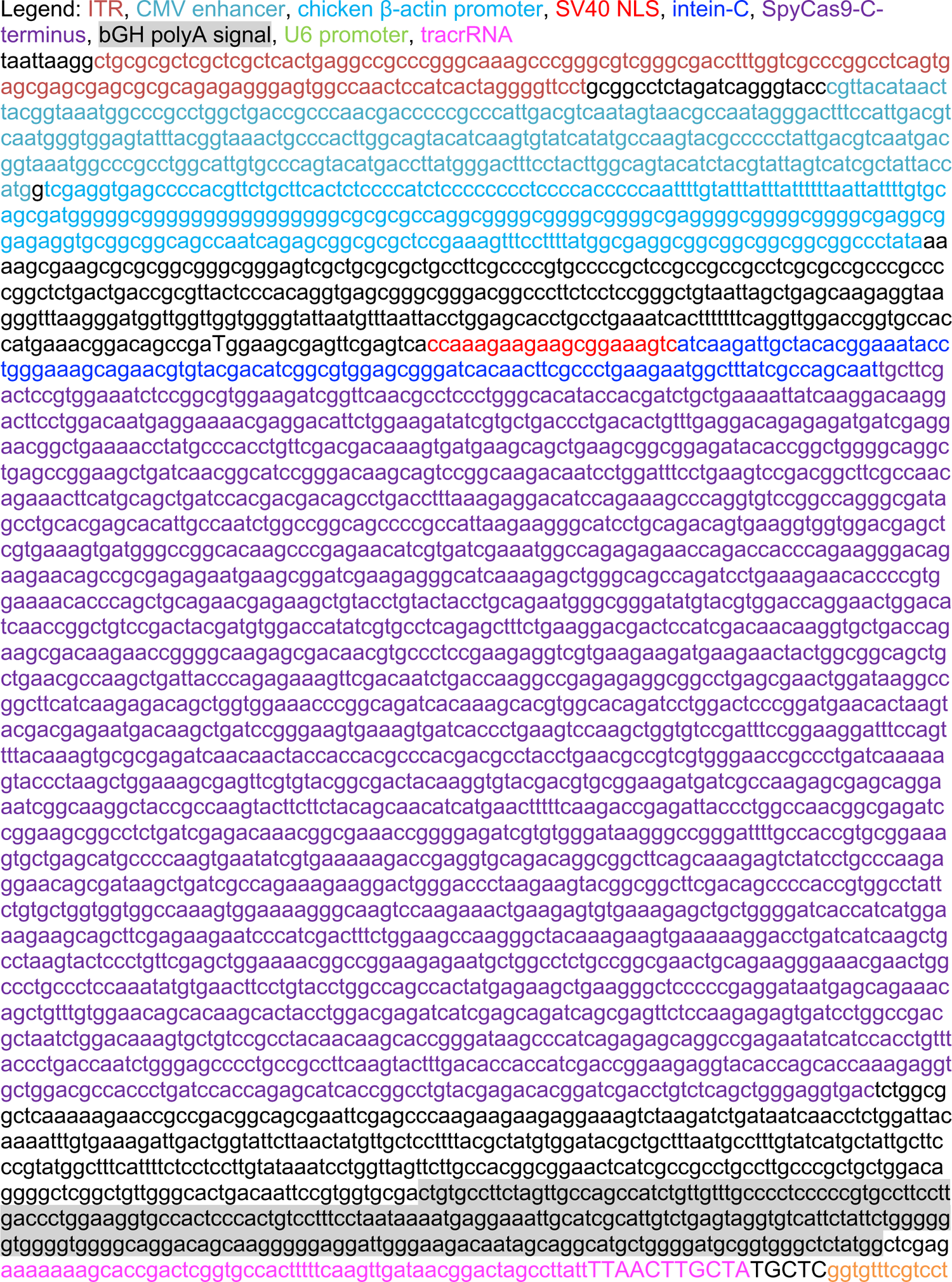
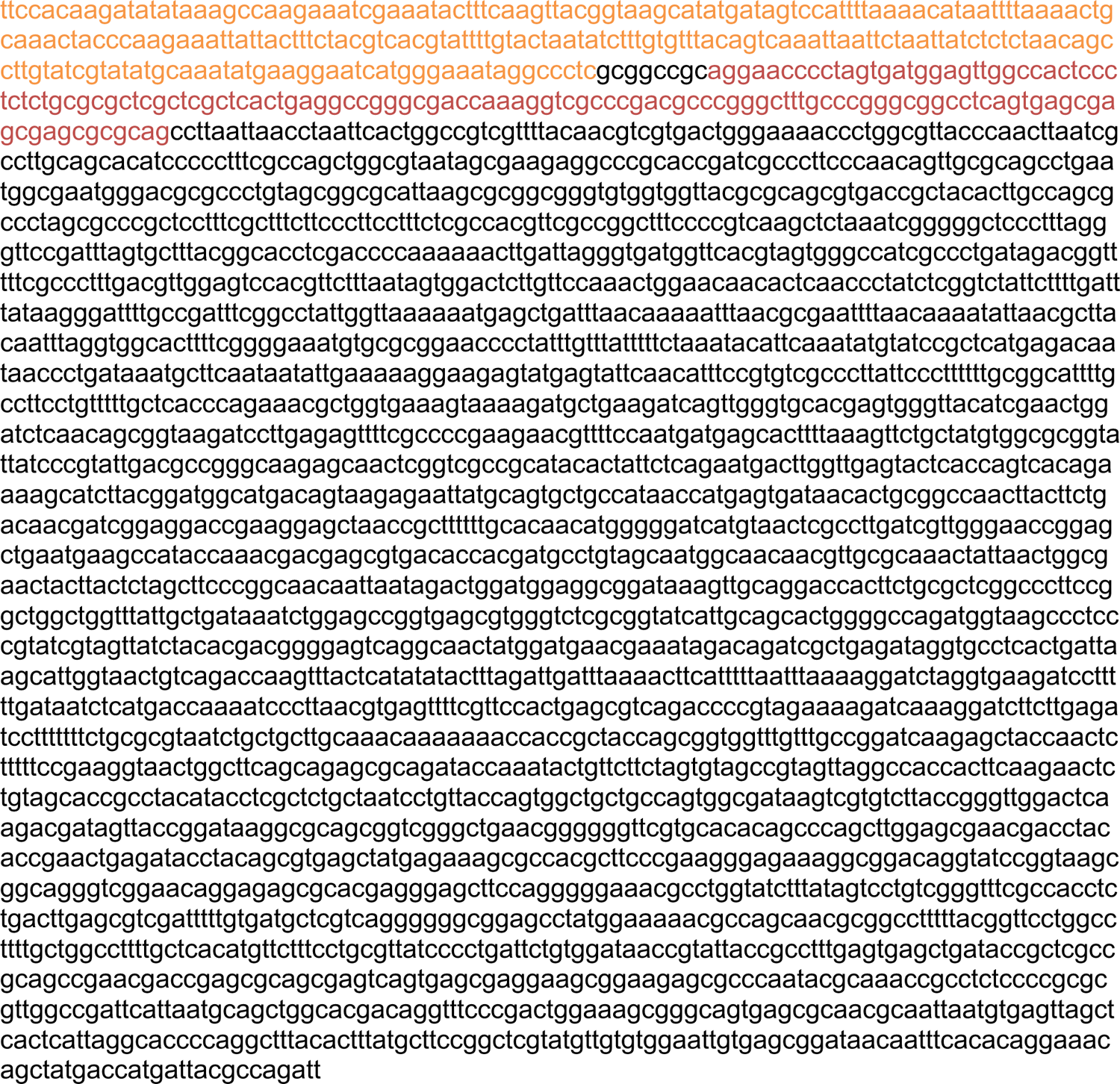

## References

1. Cox, D.B.T., Platt, R.J. and Zhang, F. (2015) Therapeutic genome editing: prospects and challenges. Nat. Med., 21, 121–131.

2. Hsu, P.D., Lander, E.S. and Zhang, F. (2014) Development and applications of CRISPR-Cas9 for genome engineering. Cell, 157, 1262–1278.

3. Anzalone, A.V., Koblan, L.W. and Liu, D.R. (2020) Genome editing with CRISPR–Cas nucleases, base editors, transposases and prime editors. Nat. Biotechnol., 38, 824–844.

4. Jinek, M., Chylinski, K., Fonfara, I., Hauer, M., Doudna, J.A. and Charpentier, E. (2012) A Programmable Dual-RNA–Guided DNA Endonuclease in Adaptive Bacterial Immunity. Science, 337, 816–821.

5. Deltcheva, E., Chylinski, K., Sharma, C.M., Gonzales, K., Chao, Y., Pirzada, Z.A., Eckert, M.R., Vogel, J. and Charpentier, E. (2011) CRISPR RNA maturation by trans-encoded small RNA and host factor RNase III. Nature, 471, 602–607.

6. Cong, L., Ran, F.A., Cox, D., Lin, S., Barretto, R., Habib, N., Hsu, P.D., Wu, X., Jiang, W., Marraffini, L.A., et al. (2013) Multiplex genome engineering using CRISPR/Cas systems. Science, 339, 819–823.

7. Doudna, J.A. and Charpentier, E. (2014) The new frontier of genome engineering with CRISPR-Cas9. Science, 346, 1258096.

8. Gaudelli, N.M., Komor, A.C., Rees, H.A., Packer, M.S., Badran, A.H., Bryson, D.I. and Liu, D.R. (2017) Programmable base editing of A•T to G•C in genomic DNA without DNA cleavage. Nature, 551, 464–471.

9. Komor, A.C., Kim, Y.B., Packer, M.S., Zuris, J.A. and Liu, D.R. (2016) Programmable editing of a target base in genomic DNA without double-stranded DNA cleavage. Nature, 533, 420– 424.

10. Koblan, L.W., Arbab, M., Shen, M.W., Hussmann, J.A., Anzalone, A.V., Doman, J.L., Newby, G.A., Yang, D., Mok, B., Replogle, J.M., et al. (2021) Efficient C•G-to-G•C base editors developed using CRISPRi screens, target-library analysis, and machine learning. Nat. Biotechnol., 39, 1414–1425.

11. Richter, M.F., Zhao, K.T., Eton, E., Lapinaite, A., Newby, G.A., Thuronyi, B.W., Wilson, C., Koblan, L.W., Zeng, J., Bauer, D.E., et al. (2020) Phage-assisted evolution of an adenine base editor with improved Cas domain compatibility and activity. Nat. Biotechnol., 38, 883– 891.

12. Rees, H.A. and Liu, D.R. (2018) Base editing: precision chemistry on the genome and transcriptome of living cells. Nat. Rev. Genet., 19, 770–788.

13. Anzalone, A.V., Randolph, P.B., Davis, J.R., Sousa, A.A., Koblan, L.W., Levy, J.M., Chen, P.J., Wilson, C., Newby, G.A., Raguram, A., et al. (2019) Search-and-replace genome editing without double-strand breaks or donor DNA. Nature, 576, 149–157.

14. Chen, P.J. and Liu, D.R. (2022) Prime editing for precise and highly versatile genome manipulation. Nat. Rev. Genet., 10.1038/s41576-022-00541-1.

15. Newby, G.A. and Liu, D.R. (2021) In vivo somatic cell base editing and prime editing. Mol. Ther., 10.1016/j.ymthe.2021.09.002.

16. Khvorova, A. and Watts, J.K. (2017) The chemical evolution of oligonucleotide therapies of clinical utility. Nat. Biotechnol., 35, 238–248.

17. Roberts, T.C., Langer, R. and Wood, M.J.A. (2020) Advances in oligonucleotide drug delivery. Nat. Rev. Drug Discov., 19, 673–694.

18. Hassler, M.R., Turanov, A.A., Alterman, J.F., Haraszti, R.A., Coles, A.H., Osborn, M.F., Echeverria, D., Nikan, M., Salomon, W.E., Roux, L., et al. (2018) Comparison of partially and fully chemically-modified siRNA in conjugate-mediated delivery in vivo. Nucleic Acids Res., 46, 2185–2196.

19. Corey, D.R. (2007) Chemical modification: the key to clinical application of RNA interference? J. Clin. Invest., 117, 3615–3622.

20. Biscans, A., Caiazzi, J., Davis, S., McHugh, N., Sousa, J. and Khvorova, A. (2020) The chemical structure and phosphorothioate content of hydrophobically modified siRNAs impact extrahepatic distribution and efficacy. Nucleic Acids Res., 48, 7665–7680.

21. Allerson, C.R., Sioufi, N., Jarres, R., Prakash, T.P., Naik, N., Berdeja, A., Wanders, L., Griffey, R.H., Swayze, E.E. and Bhat, B. (2005) Fully 2’-modified oligonucleotide duplexes with improved in vitro potency and stability compared to unmodified small interfering RNA. J. Med. Chem., 48, 901–904.

22. Dowdy, S.F. (2017) Overcoming cellular barriers for RNA therapeutics. Nat. Biotechnol., 35, 222–229.

23. Mir, A., Alterman, J.F., Hassler, M.R., Debacker, A.J., Hudgens, E., Echeverria, D., Brodsky, M.H., Khvorova, A., Watts, J.K. and Sontheimer, E.J. (2018) Heavily and fully modified RNAs guide efficient SpyCas9-mediated genome editing. Nat. Commun., 9, 2641.

24. Hendel, A., Bak, R.O., Clark, J.T., Kennedy, A.B., Ryan, D.E., Roy, S., Steinfeld, I., Lunstad, B.D., Kaiser, R.J., Wilkens, A.B., et al. (2015) Chemically modified guide RNAs enhance CRISPR-Cas genome editing in human primary cells. Nat. Biotechnol., 33, 985– 989.

25. Rahdar, M., McMahon, M.A., Prakash, T.P., Swayze, E.E., Bennett, C.F. and Cleveland, D.W. (2015) Synthetic CRISPR RNA-Cas9–guided genome editing in human cells. Proceedings of the National Academy of Sciences, 112, E7110–E7117.

26. Ryan, D.E., Taussig, D., Steinfeld, I., Phadnis, S.M., Lunstad, B.D., Singh, M., Vuong, X., Okochi, K.D., McCaffrey, R., Olesiak, M., et al. (2017) Improving CRISPR–Cas specificity with chemical modifications in single-guide RNAs. Nucleic Acids Res., 46, 792–803.

27. Rozners, E. (2022) Chemical Modifications of CRISPR RNAs to Improve Gene-Editing Activity and Specificity. J. Am. Chem. Soc., 144, 12584–12594.

28. Bost, J.P., Barriga, H., Holme, M.N., Gallud, A., Maugeri, M., Gupta, D., Lehto, T., Valadi, H., Esbjörner, E.K., Stevens, M.M., et al. (2021) Delivery of Oligonucleotide Therapeutics: Chemical Modifications, Lipid Nanoparticles, and Extracellular Vesicles. ACS Nano, 15, 13993–14021.

29. Chernikov, I.V., Vlassov, V.V. and Chernolovskaya, E.L. (2019) Current Development of siRNA Bioconjugates: From Research to the Clinic. Front. Pharmacol., 10, 444.

30. Gangopadhyay, S. and Gore, K.R. (2022) Advances in siRNA therapeutics and synergistic effect on siRNA activity using emerging dual ribose modifications. RNA Biol., 19, 452–467.

31. Shen, X. and Corey, D.R. (2018) Chemistry, mechanism and clinical status of antisense oligonucleotides and duplex RNAs. Nucleic Acids Res., 46, 1584–1600.

32. Traber, G.M. and Yu, A.-M. (2022) RNAi Based Therapeutics and Novel RNA Bioengineering Technologies. J. Pharmacol. Exp. Ther., 10.1124/jpet.122.001234.

33. Humphreys, S.C., Davis, J.A., Iqbal, S., Kamel, A., Kulmatycki, K., Lao, Y., Liu, X., Rodgers, J., Snoeys, J., Vigil, A., et al. (2022) Considerations and recommendations for assessment of plasma protein binding and drug–drug interactions for siRNA therapeutics. Nucleic Acids Res., 50, 6020–6037.

34. Jafar-Nejad, P., Powers, B., Soriano, A., Zhao, H., Norris, D.A., Matson, J., DeBrosse-Serra, B., Watson, J., Narayanan, P., Chun, S.J., et al. (2021) The atlas of RNase H antisense oligonucleotide distribution and activity in the CNS of rodents and non-human primates following central administration. Nucleic Acids Res., 49, 657–673.

35. Alterman, J.F., Godinho, B.M.D.C., Hassler, M.R., Ferguson, C.M., Echeverria, D., Sapp, E., Haraszti, R.A., Coles, A.H., Conroy, F., Miller, R., et al. (2019) A divalent siRNA chemical scaffold for potent and sustained modulation of gene expression throughout the central nervous system. Nat. Biotechnol., 37, 884–894.

36. Kuzmin, D.A., Shutova, M.V., Johnston, N.R., Smith, O.P., Fedorin, V.V., Kukushkin, Y.S., van der Loo, J.C.M. and Johnstone, E.C. (2021) The clinical landscape for AAV gene therapies. Nature Reviews Drug Discovery, 20, 173–174.

37. Wang, D., Tai, P.W.L. and Gao, G. (2019) Adeno-associated virus vector as a platform for gene therapy delivery. Nat. Rev. Drug Discov., 18, 358–378.

38. Raguram, A., Banskota, S. and Liu, D.R. (2022) Therapeutic in vivo delivery of gene editing agents. Cell, 185, 2806–2827.

39. Hocquemiller, M., Giersch, L., Audrain, M., Parker, S. and Cartier, N. (2016) Adeno-Associated Virus-Based Gene Therapy for CNS Diseases. Hum. Gene Ther., 27, 478–496.

40. Iyer, S., Mir, A., Vega-Badillo, J., Roscoe, B.P., Ibraheim, R., Zhu, L.J., Lee, J., Liu, P., Luk, K., Mintzer, E., et al. (2022) Efficient Homology-Directed Repair with Circular Single-Stranded DNA Donors. CRISPR J, 5, 685–701.

41. Sena-Esteves, M. and Gao, G. (2020) Introducing Genes into Mammalian Cells: Viral Vectors. Cold Spring Harb. Protoc., 2020, 095513.

42. Zhang, H., Bamidele, N., Liu, P., Ojelabi, O., Gao, X.D., Rodriguez, T., Cheng, H., Kelly, K., Watts, J.K., Xie, J., et al. (2022) Adenine Base Editing In Vivo with a Single Adeno-Associated Virus Vector. GEN Biotechnol, 1, 285–299.

43. Carroll, J.B., Warby, S.C., Southwell, A.L., Doty, C.N., Greenlee, S., Skotte, N., Hung, G., Frank Bennett, C., Freier, S.M. and Hayden, M.R. (2011) Potent and Selective Antisense Oligonucleotides Targeting Single-Nucleotide Polymorphisms in the Huntington Disease Gene / Allele-Specific Silencing of Mutant Huntingtin. Molecular Therapy, 19, 2178–2185.

44. Song, C.-Q., Jiang, T., Richter, M., Rhym, L.H., Koblan, L.W., Zafra, M.P., Schatoff, E.M., Doman, J.L., Cao, Y., Dow, L.E., et al. (2020) Adenine base editing in an adult mouse model of tyrosinaemia. Nat Biomed Eng, 4, 125–130.

45. Clement, K., Rees, H., Canver, M.C., Gehrke, J.M., Farouni, R., Hsu, J.Y., Cole, M.A., Liu, D.R., Joung, J.K., Bauer, D.E., et al. (2019) CRISPResso2 provides accurate and rapid genome editing sequence analysis. Nat. Biotechnol., 37, 224–226.

46. Certo, M.T., Ryu, B.Y., Annis, J.E., Garibov, M., Jarjour, J., Rawlings, D.J. and Scharenberg, A.M. (2011) Tracking genome engineering outcome at individual DNA breakpoints. Nature Methods, 8, 671–676.

47. Sontheimer, E. J., Khvorova, A., Watts, J. K., Amrani, N., Chen, Z., Hassler, M., Moreno, D. E., Alterman, J.F., Wolfe, S.A., Yamada, K., Devi, G., Zhang, H. (2021) Modified guide rnas for crispr genome editing. U.S. Patent Application No*. 17/318,* 846.

48. Nishina, K., Piao, W., Yoshida-Tanaka, K., Sujino, Y., Nishina, T., Yamamoto, T., Nitta, K., Yoshioka, K., Kuwahara, H., Yasuhara, H., et al. (2015) DNA/RNA heteroduplex oligonucleotide for highly efficient gene silencing. Nat. Commun., 6, 7969.

49. Nagata, T., Dwyer, C.A., Yoshida-Tanaka, K., Ihara, K., Ohyagi, M., Kaburagi, H., Miyata, H., Ebihara, S., Yoshioka, K., Ishii, T., et al. (2021) Cholesterol-functionalized DNA/RNA heteroduplexes cross the blood–brain barrier and knock down genes in the rodent CNS. Nat. Biotechnol., 39, 1529–1536.

50. Allison, S.J. and Milner, J. (2014) RNA Interference by Single- and Double-stranded siRNA With a DNA Extension Containing a 3′ Nuclease-resistant Mini-hairpin Structure. Molecular Therapy - Nucleic Acids, 3, e141.

51. Xu, Y., Linde, A., Larsson, O., Thormeyer, D., Elmen, J., Wahlestedt, C. and Liang, Z. (2004) Functional comparison of single- and double-stranded siRNAs in mammalian cells. Biochem. Biophys. Res. Commun., 316, 680–687.

52. Zhang, K., Hodge, J., Chatterjee, A., Moon, T.S. and Parker, K.M. (2021) Duplex Structure of Double-Stranded RNA Provides Stability against Hydrolysis Relative to Single-Stranded RNA. Environ. Sci. Technol., 55, 8045–8053.

53. Gehl, J. (2003) Electroporation: theory and methods, perspectives for drug delivery, gene therapy and research. Acta Physiol. Scand., 177, 437–447.

54. Alterman, J.F., Hall, L.M., Coles, A.H., Hassler, M.R., Didiot, M.-C., Chase, K., Abraham, J., Sottosanti, E., Johnson, E., Sapp, E., et al. (2015) Hydrophobically Modified siRNAs Silence Huntingtin mRNA in Primary Neurons and Mouse Brain. Mol. Ther. Nucleic Acids, 4, e266.

55. Linnane, E., Davey, P., Zhang, P., Puri, S., Edbrooke, M., Chiarparin, E., Revenko, A.S., Macleod, A.R., Norman, J.C. and Ross, S.J. (2019) Differential uptake, kinetics and mechanisms of intracellular trafficking of next-generation antisense oligonucleotides across human cancer cell lines. Nucleic Acids Res., 47, 4375–4392.

56. Wang, S., Allen, N., Vickers, T.A., Revenko, A.S., Sun, H., Liang, X.-H. and Crooke, S.T. (2018) Cellular uptake mediated by epidermal growth factor receptor facilitates the intracellular activity of phosphorothioate-modified antisense oligonucleotides. Nucleic Acids Res., 46, 3579–3594.

57. Benizri, S., Gissot, A., Martin, A., Vialet, B., Grinstaff, M.W. and Barthélémy, P. (2019) Bioconjugated Oligonucleotides: Recent Developments and Therapeutic Applications. Bioconjug. Chem., 30, 366–383.

58. Springer, A.D. and Dowdy, S.F. (2018) GalNAc-siRNA Conjugates: Leading the Way for Delivery of RNAi Therapeutics. Nucleic Acid Ther., 28, 109–118.

59. Debacker, A.J., Voutila, J., Catley, M., Blakey, D. and Habib, N. (2020) Delivery of Oligonucleotides to the Liver with GalNAc: From Research to Registered Therapeutic Drug. Mol. Ther., 28, 1759–1771.

60. Osborn, M.F. and Khvorova, A. (2018) Improving siRNA Delivery In Vivo Through Lipid Conjugation. Nucleic Acid Ther., 28, 128–136.

61. Biscans, A., Coles, A., Haraszti, R., Echeverria, D., Hassler, M., Osborn, M. and Khvorova, A. (2019) Diverse lipid conjugates for functional extra-hepatic siRNA delivery in vivo. Nucleic Acids Res., 47, 1082–1096.

62. Nikan, M., Osborn, M.F., Coles, A.H., Godinho, B.M., Hall, L.M., Haraszti, R.A., Hassler, M.R., Echeverria, D., Aronin, N. and Khvorova, A. (2016) Docosahexaenoic Acid Conjugation Enhances Distribution and Safety of siRNA upon Local Administration in Mouse Brain. Mol. Ther. Nucleic Acids, 5, e344.

63. Biscans, A., Caiazzi, J., McHugh, N., Hariharan, V., Muhuri, M. and Khvorova, A. (2021) Docosanoic acid conjugation to siRNA enables functional and safe delivery to skeletal and cardiac muscles. Mol. Ther., 29, 1382–1394.

64. Soutschek, J., Akinc, A., Bramlage, B., Charisse, K., Constien, R., Donoghue, M., Elbashir, S., Geick, A., Hadwiger, P., Harborth, J., et al. (2004) Therapeutic silencing of an endogenous gene by systemic administration of modified siRNAs. Nature, 432, 173–178.

65. Brown, K.M., Nair, J.K., Janas, M.M., Anglero-Rodriguez, Y.I., Dang, L.T.H., Peng, H., Theile, C.S., Castellanos-Rizaldos, E., Brown, C., Foster, D., et al. (2022) Expanding RNAi therapeutics to extrahepatic tissues with lipophilic conjugates. Nat. Biotechnol., 40, 1500– 1508.

66. Biscans, A., Coles, A., Echeverria, D. and Khvorova, A. (2019) The valency of fatty acid conjugates impacts siRNA pharmacokinetics, distribution, and efficacy in vivo. J. Control. Release, 302, 116–125.

67. Østergaard, M.E., Jackson, M., Low, A., E Chappell, A., G Lee, R., Peralta, R.Q., Yu, J., Kinberger, G.A., Dan, A., Carty, R., et al. (2019) Conjugation of hydrophobic moieties enhances potency of antisense oligonucleotides in the muscle of rodents and non-human primates. Nucleic Acids Res., 47, 6045–6058.

68. Tran, P., Weldemichael, T., Liu, Z. and Li, H.-Y. (2022) Delivery of Oligonucleotides: Efficiency with Lipid Conjugation and Clinical Outcome. Pharmaceutics, 14.

69. Nishina, K., Unno, T., Uno, Y., Kubodera, T., Kanouchi, T., Mizusawa, H. and Yokota, T. (2008) Efficient In Vivo Delivery of siRNA to the Liver by Conjugation of α-Tocopherol. Molecular Therapy, 10.1038/sj.mt.6300420.

70. Nishina, T., Numata, J., Nishina, K., Yoshida-Tanaka, K., Nitta, K., Piao, W., Iwata, R., Ito, S., Kuwahara, H., Wada, T., et al. (2015) Chimeric Antisense Oligonucleotide Conjugated to α-Tocopherol. Molecular Therapy - Nucleic Acids, 4, e220.

71. Lau, S., Graham, B., Cao, N., Boyd, B.J., Pouton, C.W. and White, P.J. (2012) Enhanced Extravasation, Stability and *in Vivo* Cardiac Gene Silencing via *in Situ* siRNA–Albumin Conjugation. Molecular Pharmaceutics, 9, 71–80.

72. Sharma, V.K., Osborn, M.F., Hassler, M.R., Echeverria, D., Ly, S., Ulashchik, E.A., Martynenko-Makaev, Y.V., Shmanai, V.V., Zatsepin, T.S., Khvorova, A., et al. (2018) Novel Cluster and Monomer-Based GalNAc Structures Induce Effective Uptake of siRNAs in Vitro and in Vivo. Bioconjug. Chem., 29, 2478–2488.

73. Aronin, N. (2006) Target selectivity in mRNA silencing. Gene Therapy, 13, 509–516.

74. Bäumer, S., Bäumer, N., Appel, N., Terheyden, L., Fremerey, J., Schelhaas, S., Wardelmann, E., Buchholz, F., Berdel, W.E. and Müller-Tidow, C. (2015) Antibody-mediated delivery of anti-KRAS-siRNA in vivo overcomes therapy resistance in colon cancer. Clin. Cancer Res., 21, 1383–1394.

75. Song, E., Zhu, P., Lee, S.-K., Chowdhury, D., Kussman, S., Dykxhoorn, D.M., Feng, Y., Palliser, D., Weiner, D.B., Shankar, P., et al. (2005) Antibody mediated in vivo delivery of small interfering RNAs via cell-surface receptors. Nat. Biotechnol., 23, 709–717.

76. Muzumdar, M.D., Tasic, B., Miyamichi, K., Li, L. and Luo, L. (2007) A global double-fluorescent Cre reporter mouse. genesis, 45, 593–605.

77. Cui, H., Zhu, X., Li, S., Wang, P. and Fang, J. (2021) Liver-Targeted Delivery of Oligonucleotides with N-Acetylgalactosamine Conjugation. ACS Omega, 6, 16259–16265.

78. Aponte, J.L., Sega, G.A., Hauser, L.J., Dhar, M.S., Withrow, C.M., Carpenter, D.A., Rinchik, E.M., Culiat, C.T. and Johnson, D.K. (2001) Point mutations in the murine fumarylacetoacetate hydrolase gene: Animal models for the human genetic disorder hereditary tyrosinemia type 1. Proceedings of the National Academy of Sciences, 98, 641– 645.

79. Lindstedt, S. (1992) Treatment of hereditary tyrosinaemia type I by inhibition of 4-hydroxyphenylpyruvate dioxygenase. The Lancet, 340, 813–817.

80. Jiang, T., Henderson, J.M., Coote, K., Cheng, Y., Valley, H.C., Zhang, X.-O., Wang, Q., Rhym, L.H., Cao, Y., Newby, G.A., et al. (2020) Chemical modifications of adenine base editor mRNA and guide RNA expand its application scope. Nat. Commun., 11, 1979.

81. Levy, J.M., Yeh, W.-H., Pendse, N., Davis, J.R., Hennessey, E., Butcher, R., Koblan, L.W., Comander, J., Liu, Q. and Liu, D.R. (2020) Cytosine and adenine base editing of the brain, liver, retina, heart and skeletal muscle of mice via adeno-associated viruses. Nat Biomed Eng, 4, 97–110.

82. Alves, C.R.R., Ha, L.L., Yaworski, R., Lazzarotto, C.R., Christie, K.A., Reilly, A., Beauvais, A., Doll, R.M., de la Cruz, D., Maguire, C.A., et al. (2023) Base editing as a genetic treatment for spinal muscular atrophy. *bioRxiv*, 10.1101/2023.01.20.524978.

83. Davis, J.R., Wang, X., Witte, I.P., Huang, T.P., Levy, J.M., Raguram, A., Banskota, S., Seidah, N.G., Musunuru, K. and Liu, D.R. (2022) Efficient in vivo base editing via single adeno-associated viruses with size-optimized genomes encoding compact adenine base editors. Nat Biomed Eng, 6, 1272–1283.

84. Schwartz, A.L., Fridovich, S.E. and Lodish, H.F. (1982) Kinetics of internalization and recycling of the asialoglycoprotein receptor in a hepatoma cell line. J. Biol. Chem., 257, 4230–4237.

85. Nair, J.K., Willoughby, J.L.S., Chan, A., Charisse, K., Alam, M.R., Wang, Q., Hoekstra, M., Kandasamy, P., Kel’in, A.V., Milstein, S., et al. (2014) Multivalent N-acetylgalactosamine-conjugated siRNA localizes in hepatocytes and elicits robust RNAi-mediated gene silencing. J. Am. Chem. Soc., 136, 16958–16961.

86. Bon, C., Hofer, T., Bousquet-Mélou, A., Davies, M.R. and Krippendorff, B.-F. (2017) Capacity limits of asialoglycoprotein receptor-mediated liver targeting. MAbs, 9, 1360–1369.

87. Crooke, S.T., Witztum, J.L., Bennett, C.F. and Baker, B.F. (2019) RNA-Targeted Therapeutics. Cell Metab., 29, 501.

88. Ohrt, T., Muetze, J., Svoboda, P. and Schwille, P. (2012) Intracellular localization and routing of miRNA and RNAi pathway components. Curr. Top. Med. Chem., 12, 79–88.

89. Godinho, B.M.D.C., Knox, E.G., Hildebrand, S., Gilbert, J.W., Echeverria, D., Kennedy, Z., Haraszti, R.A., Ferguson, C.M., Coles, A.H., Biscans, A., et al. (2022) PK-modifying anchors significantly alter clearance kinetics, tissue distribution, and efficacy of therapeutics siRNAs. Mol. Ther. Nucleic Acids, 29, 116–132.

90. Gonzalez, T.J., Simon, K.E., Blondel, L.O., Fanous, M.M., Roger, A.L., Maysonet, M.S., Devlin, G.W., Smith, T.J., Oh, D.K., Havlik, L.P., et al. (2022) Cross-species evolution of a highly potent AAV variant for therapeutic gene transfer and genome editing. Nat. Commun., 13, 5947.

91. Goertsen, D., Flytzanis, N.C., Goeden, N., Chuapoco, M.R., Cummins, A., Chen, Y., Fan, Y., Zhang, Q., Sharma, J., Duan, Y., et al. (2022) AAV capsid variants with brain-wide transgene expression and decreased liver targeting after intravenous delivery in mouse and marmoset. Nat. Neurosci., 25, 106–115.

92. Hudry, E. and Vandenberghe, L.H. (2019) Therapeutic AAV Gene Transfer to the Nervous System: A Clinical Reality. Neuron, 102, 263.

93. Godinho, B.M.D.C., Henninger, N., Bouley, J., Alterman, J.F., Haraszti, R.A., Gilbert, J.W., Sapp, E., Coles, A.H., Biscans, A., Nikan, M., et al. (2018) Transvascular Delivery of Hydrophobically Modified siRNAs: Gene Silencing in the Rat Brain upon Disruption of the Blood-Brain Barrier. Mol. Ther., 26, 2580–2591.

94. Ferguson, C.M., Godinho, B.M., Alterman, J.F., Coles, A.H., Hassler, M., Echeverria, D., Gilbert, J.W., Knox, E.G., Caiazzi, J., Haraszti, R.A., et al. (2021) Comparative route of administration studies using therapeutic siRNAs show widespread gene modulation in Dorset sheep. JCI Insight, 6.

95. Fischell, J.M. and Fishman, P.S. (2021) A Multifaceted Approach to Optimizing AAV Delivery to the Brain for the Treatment of Neurodegenerative Diseases. Front. Neurosci., 15, 747726.

96. Barua, N.U., Gill, S.S. and Love, S. (2014) Convection-enhanced drug delivery to the brain: therapeutic potential and neuropathological considerations. Brain Pathol., 24, 117–127.

